# Identification of rare transient somatic cell states induced by injury and required for whole-body regeneration

**DOI:** 10.1101/2020.06.04.132753

**Authors:** Blair W. Benham-Pyle, Carolyn E. Brewster, Aubrey M. Kent, Frederick G. Mann, Shiyuan Chen, Allison R. Scott, Andrew C. Box, Alejandro Sánchez Alvarado

**Affiliations:** Stowers Institute for Medical Research, Kansas City, MO, USA; Howard Hughes Institute for Medical Research, Kansas City, MO, USA

## Abstract

Regeneration requires functional coordination of stem cells, their progeny, and differentiated cells. Past studies have focused on regulation of stem cell identity and proliferation near to the wound-site, but less is known about contributions made by differentiated cells distant to the injury. Here, we present a comprehensive atlas of whole-body regeneration over time and identify rare, transient, somatic cell states induced by injury and required for regeneration. To characterize amputation-specific signaling across a whole animal, 299,998 single-cell transcriptomes were captured from planarian tissue fragments competent and incompetent to regenerate. Amputation-specific cell states were rare, non-uniformly distributed across tissues, and particularly enriched in muscle (mesoderm), epidermis (ectoderm), and intestine (endoderm). Moreover, RNAi-mediated knockdown of genes up-regulated in amputation-specific cell states drastically reduced regenerative capacity. These results identify novel cell states and molecules required for whole-body regeneration and indicate that regenerative capacity requires transcriptional plasticity in a rare subset of differentiated cells.

## Main Text

Robust injury and tissue repair mechanisms provide fitness advantages to organisms, but regenerative capacity is heterogeneously distributed across the animal kingdom. Numerous invertebrate species have the capacity to regenerate an entire organism from dissociated cells (*1*) or small tissue fragments (*2–7*). In vertebrates, both teleost fish and urodele amphibians have the capacity to regenerate appendages and organs after damage or amputation (*8–12*). Even in mammals, which have relatively limited regenerative potential, numerous organ systems are capable of replacing damaged tissue or re-populating ablated tissue compartments (*13–16*). The repair of tissue damaged by injury, aging, or disease requires coordinated signaling between differentiated and proliferating cells. In adult tissues, growth is achieved by dedicated resident stem cells maintained by multicellular niches (*17–20*). Injury results in an increase in functional plasticity in both niche cells and resident stem cells, which facilitates the replacement of missing cell types and the re-establishment of tissue homeostasis. During whole-body regeneration, pluripotent adult stem cells mobilize to sites of injury, proliferate, and differentiate in accordance with local patterning cues to replace missing tissues (*3, 6, 21–24*). Mechanisms coordinating stem cell proliferation and differentiation near the wound site during whole-body regeneration have been described (*3, 22, 25, 26*). However, significantly less is known about the transcriptional states that occur in pre-existing differentiated tissues or how they support regeneration.

The free-living planarian *Schmidtea mediterranea* provides a unique opportunity to study functional plasticity across an entire animal during whole-body regeneration. Planaria have an extraordinary ability to repair or regenerate any organ system, and restore proper body proportion and tissue composition from tiny body fragments (*2*). It was recently shown that asexual *S. mediterranea* produce progeny of a fixed size (~1.2mm long) by transverse fission, independent of parent length (*27*), indicating that there may be an optimally sized tissue fragment for robust regeneration. T.H Morgan first showed that a tissue fragment amounting to 1/279^th^ of the animal from which it was taken could regenerate all missing tissues (*28*). Since then, surface area to volume models have been used to estimate that regeneration requires fragments containing ~10,000 cells (*29, 30*). The small number of cells required for regeneration, combined with robust methods for single cell transcriptomics and RNAi, facilitates characterization of transcriptional states required for whole-body regeneration. Here, we report a comprehensive atlas of successful and un-successful whole-body regeneration. We also identify and functionally test rare transient somatic cell states induced by injury and important for regenerative capacity.

### A single cell reconstruction of whole-body regeneration

We first determined the smallest *S. mediterranea* tissue fragment size competent to regenerate. Tissue biopsies of a fixed diameter ranging from 0.50mm – 1.25mm were taken from the posterior tail region of large animals. Both the survival of the resulting biopsies and regeneration of photoreceptor pigmentation were monitored following amputation (Fig. 1A).

**Figure 1.**
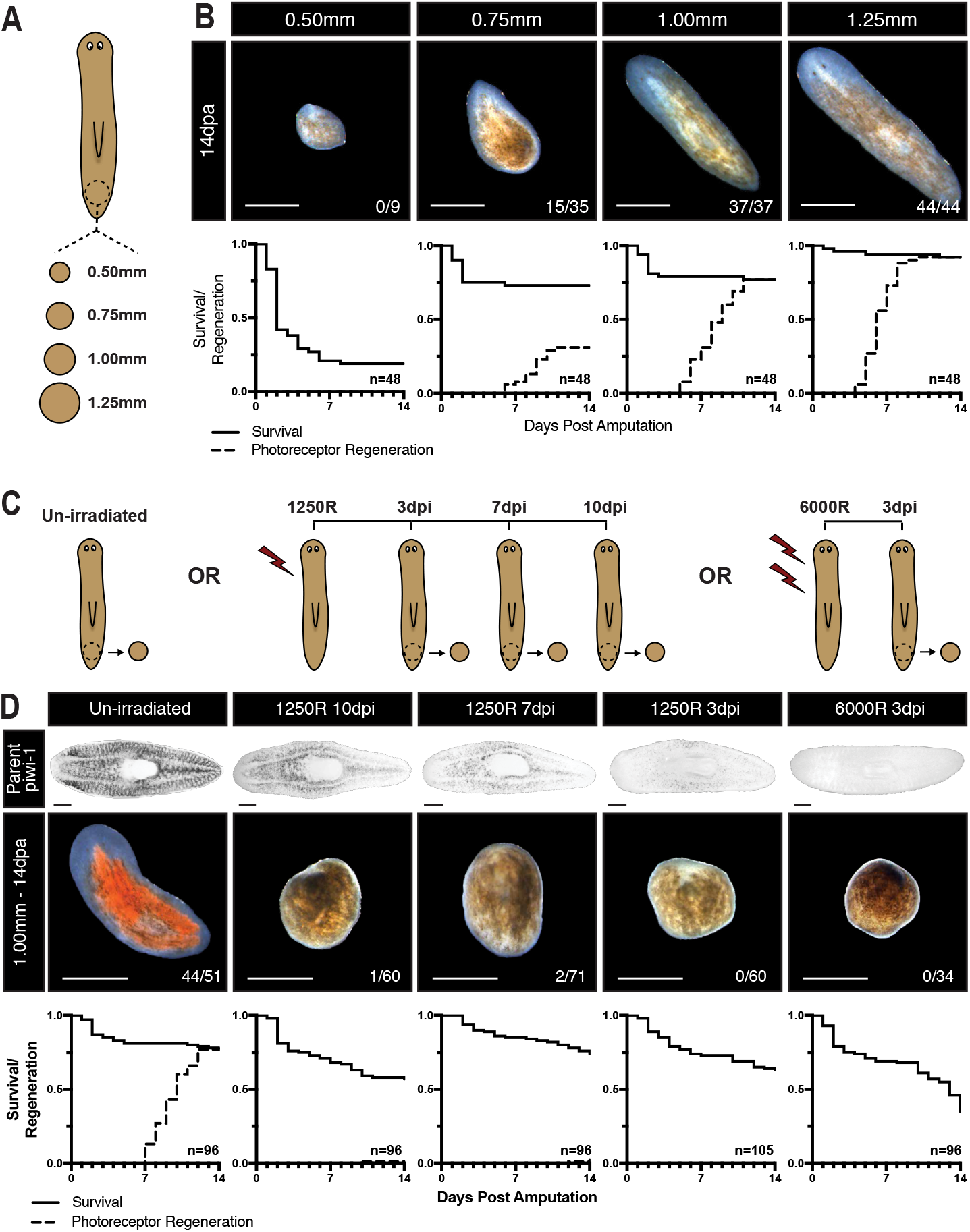
A threshold number of stem and somatic cells are required for planarian regeneration. (A) Schematic of biopsy size experiment. (B) Representative images with ratio of regenerated biopsies, survival, and photoreceptor regeneration of biopsies 0.50mm – 1.25mm in diameter 14 days after amputation. (C) Schematic of stem cell depletion biopsy experiment. (D) *piwi-1* content of unirradiated, lethally irradiated, sub-lethally irradiated animals (top) and representative morphology, survival, and photoreceptor regeneration of 1.00mm biopsies taken from those animals 14 days after amputation. Scale = 500um.

In biopsies 1.00mm in diameter and larger, all surviving tissue fragments regenerated photoreceptor pigmentation within 14 days. However, biopsies 0.75mm in diameter had reduced regenerative capacity, while biopsies 0.50mm in diameter displayed reduced survival and no ability to regenerate missing tissue (Fig. 1B). To determine whether the size-dependence of regenerative capacity was due to the number of stem cells captured in the biopsy, biopsies 0.75mm – 1.50mm in diameter were taken from animals previously treated with lethal or sub-lethal doses of ionizing radiation (Fig. 1C, Extended Data Fig.1A). Following sub-lethal irradiation, biopsies were taken at 3, 7, and 10 days post treatment, thus producing tissue fragments containing increasing numbers of *piwi-1*^+^ stem cells (Fig. 1D, top, Extended Data Fig. 1). Surprisingly, only the biopsies taken from un-irradiated animals were competent to regenerate (Fig. 1D). Indeed, the probability of successfully regenerating new pigmented photoreceptors was more dependent on the size of the tissue fragment than on the time since sub-lethal irradiation (Extended Data Fig. 1B). Altogether, these data indicate that regeneration requires a healthy tissue fragment 1.00mm in diameter (~10,000 cells) and that any damage to the stem cell compartment prior to injury dramatically reduces regenerative capacity.

In order to investigate why healthy 1.00mm biopsies were regeneration competent, while others were not, we characterized the composition and plasticity of cell states during successful and unsuccessful regeneration using single-cell RNA sequencing. Split-pool ligation-based single-cell RNA sequencing (SPLiTseq) (*31*) was used to capture single-cell transcriptomes from 1.00mm biopsies taken from un-irradiated, sub-lethally irradiated, and lethally irradiated animals (Fig.2A). Fluorescently tagged linker molecules and image cytometry were used to visualize SPLiTseq barcoding reagents in dissociated planarian cells (Extended Data Fig. 2A-B) and find optimal dye-based sort conditions to ensure that only intact barcoded cells were sequenced (Extended Data Fig. 2C-F). In order to maximize the probability of capturing rare or transient cell states, we targeted ~10,000 single-cell transcriptomes from each sample. Biopsies were dissociated, barcoded, and sequenced 0, 1, 2, 4, 7, 10, and 14 days post amputation (dpa), resulting in twenty-one samples from three regeneration time courses (Extended Data Fig. 3A-C). After filtering for transcriptome quality, the final dataset contained 299,998 single-cell transcriptomes with an average of 14,285 transcriptomes per sampled condition, 1,981 UMIs/cell, and 429 genes/cell (Fig. 2B, Extended Data Fig. 3B,C). Clustering of all captured single-cell transcriptomes identified 89 global clusters, most of which could be assigned to known tissue classes based on highly enriched genes previously reported in planarian cell type atlases and computational transfer of tissue annotations from other datasets (Fig. 2B-D, Extended Data Fig. 4-7) (*32, 33*). Once assigned to known tissues or the stem cell compartment, cell states were split into tissue-level data subsets and re-analyzed to identify additional sub-cluster diversity (Extended Data Fig. 4-7). This analysis produced tissue subclusters representing 211 distinct transcriptional states that occurred after amputation in regeneration-competent and -incompetent fragments.

**Figure 2.**
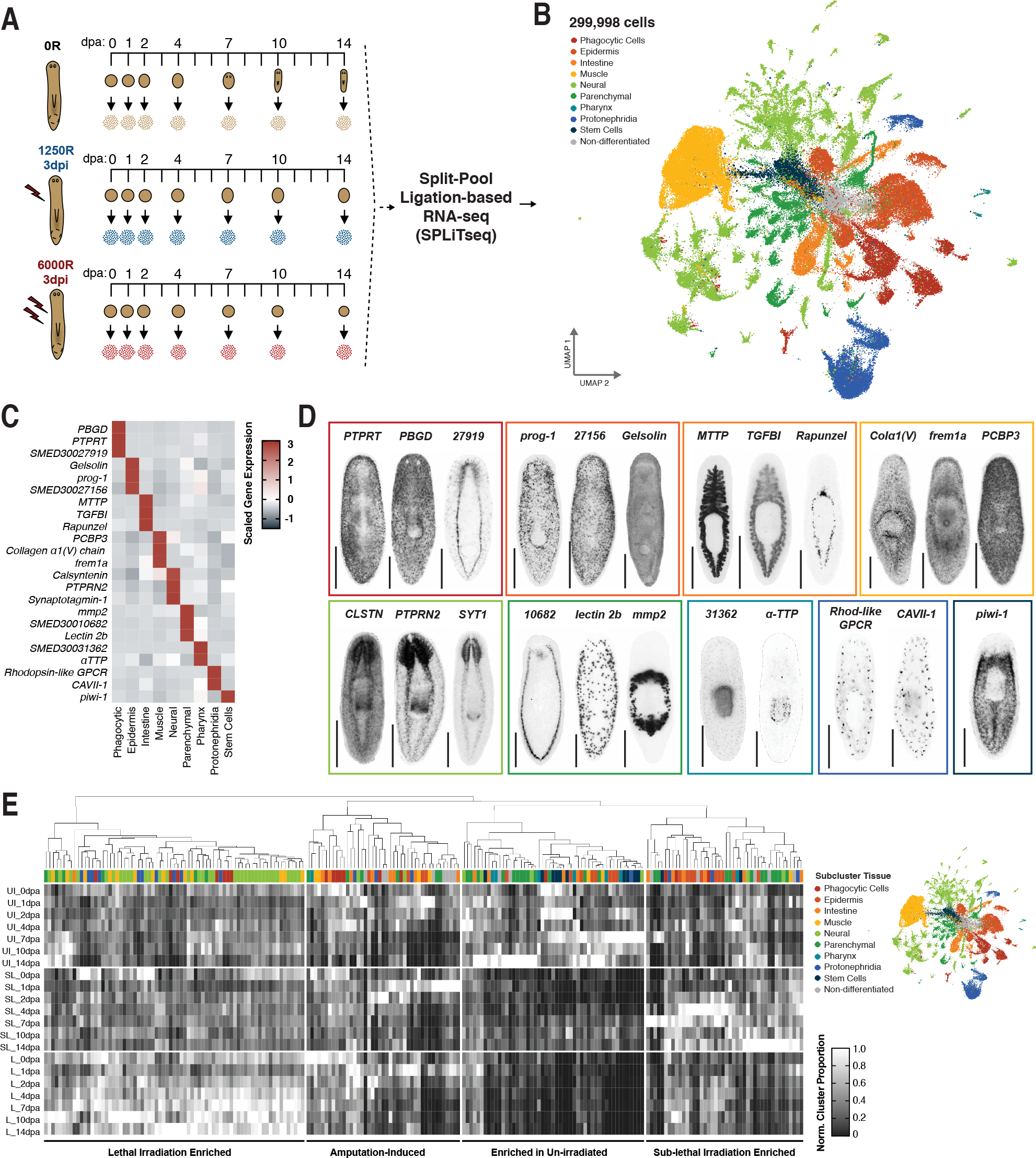
A single-cell reconstruction of successful and unsuccessful planarian regeneration. (A) Schematic depicting experimental design of single cell reconstruction. (B) UMAP embedding of captured single cell transcriptomes, colored by tissue annotation. Scaled mean gene expression in single-cell reconstruction (C) and gene expression patterns by whole mount in situ hybridization (D) of select tissue-specific markers (See also Extended Data 4-7 for additional markers and tissue-subcluster specificity). (E) Scaled proportion of cells from each tissue sub-clusters in sampled conditions, normalized to sample in which sub-cluster had maximum representation. Scale = 500um.

### Irradiation-sensitive cell states include putative stem cell states

To identify cellular components of the planarian tissue fragments contributing to successful regeneration, we first focused on cell states that only occurred in biopsies taken from un-irradiated animals. The relative contribution to animal composition made by all tissue subclusters was compared across the three regeneration time courses (Fig. 2E). When hierarchically ordered based on shared trends over the samples, cell states sorted into 4 groups – enriched in one of the three regeneration time courses or activated after amputation. Perhaps not surprisingly, numerous stem cell clusters marked by their high expression of the planarian neoblast marker *piwi-1 (34*) were among the cell states enriched in biopsies taken from unirradiated animals (Fig. 3A-C). The relative contribution of each stem cell cluster to total tissue composition confirmed that these cell states were absent in biopsies taken from lethally irradiated animals and recovered by 14 dpa in biopsies taken from sub-lethally irradiated animals (Fig. 3A-C). Genes enriched in the stem cell compartment included several known regulators of stem cell maintenance (*vasa-1, piwi-2, piwi-3, PABP1-B, MEX3b*) (*34–39*). In addition, heat shock proteins and components of the protein translation machinery (Fig. 3D, E) were also enriched in putative stem cells. RNAi-mediated knockdown (Fig. 3F) was used to test if genes enriched in irradiation sensitive clusters were required for tissue homeostasis, as would be expected for stem-cell specific genes. Following RNAi, 85% of genes enriched in irradiation-sensitive cell states produced lesions or lysis in animals at homeostasis or in regenerating tissue fragments (Figure 3G-I). These results validated the atlas and analysis approaches. Relative contribution to tissue composition across successful and unsuccessful regeneration identified cell states previously known to be required for tissue maintenance and tissue regeneration, *i.e*., putative stem cell states. Moreover, RNAi-mediated knockdown of enriched genes identified both known and novel regulators of stem cell function. Individual heat shock (*40*) and ribosomal genes (*41*) have been implicated in stem cell function during tissue maintenance and regeneration. However, our results expand upon these observations and indicate that components of the heat shock signaling pathways and protein translation machinery are required for stem cell function and tissue maintenance.

**Figure 3.**
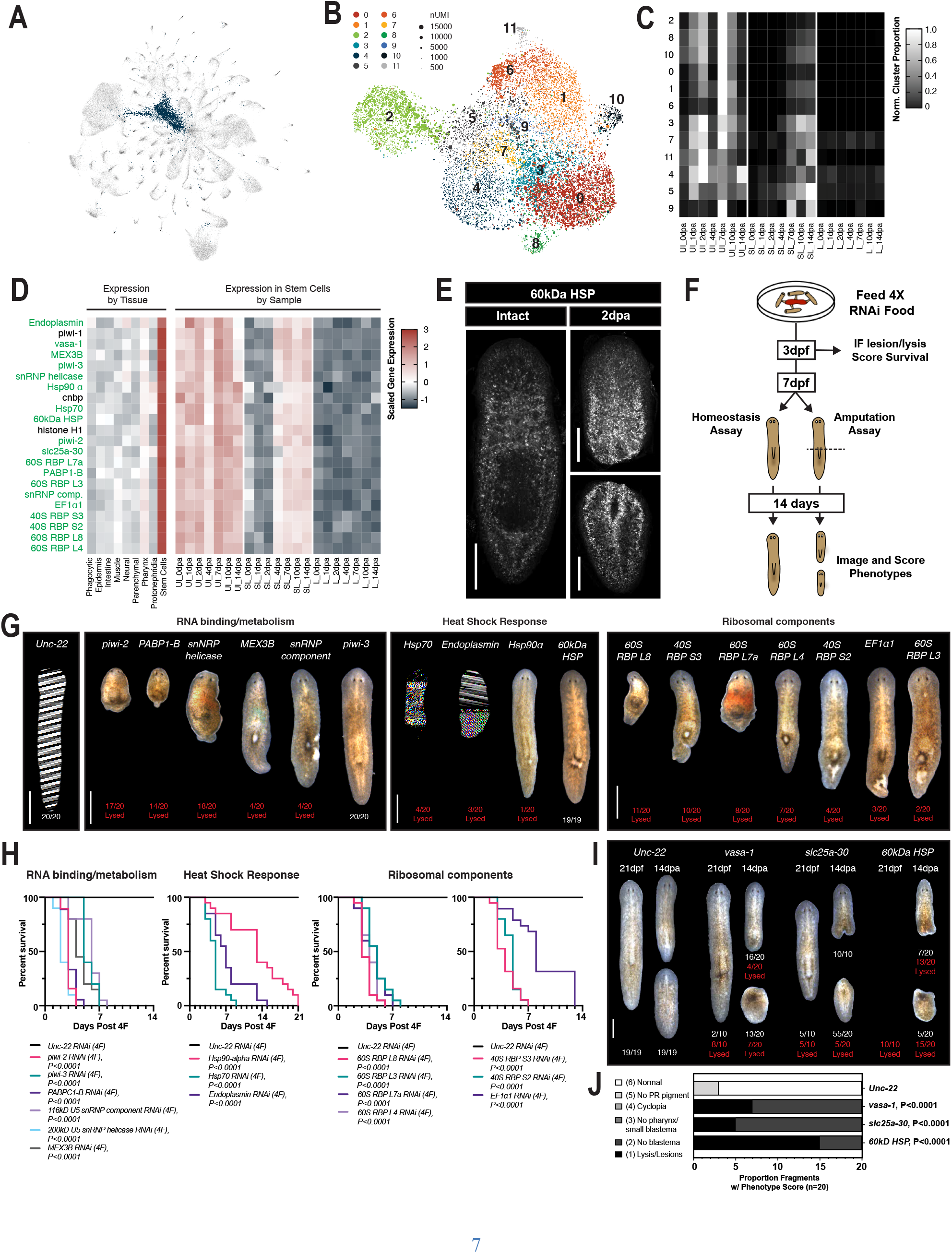
Known and novel regulators of tissue homeostasis were identified in irradiation sensitive stem cell clusters. (A) UMAP embedding of global dataset with stem cells highlighted. (B) UMAP embedding of all stem cells, colored by time after amputation. (C) Scaled proportion of cells from each stem cell sub-cluster across sampled conditions, normalized to sample in which sub-cluster had maximum representation. (D) Scaled mean expression of stem cell enriched genes by tissue and sample. (E) Z projection of confocal stack of stem cell enriched gene, *60kD HSP*. (F) Schematic representation of RNAi screen design (G) Representative images of RNAi treated animals 3 days post feeding. (H) Survival of RNAi treated animals shown in F (n=19 or 20 as noted in G). Representative images (I) of homeostatic (21 days post feeding, n=10) and regeneration phenotypes (14 days post amputation, n=20) and scoring of phenotypes (J) in RNAi treated animals. Scale = 500um. P values are Rank Log test (H) and Mann Whitney Test (J) compared to Unc-22 Control. Scale = 500um.

### Rare amputation-specific cell states are non-uniformly distributed across tissues

Given that the stem cell compartment recovered by 14 dpa in biopsies taken from sub-lethally irradiated animals (Fig. 4B) and that this recovery was not sufficient for successful regeneration (Extended Data Fig. 1B), we next focused on amputation-specific transcriptional states outside of the stem cell compartment. We expected that transcriptional states responding to a regenerative challenge would be present primarily at early time points during regeneration (24 – 96 hours after amputation). Therefore, the proportion of cells originating from each time point after amputation was calculated for all sub-clusters within known tissue subsets (Fig. 4D-K). Because several wound-induced genes important for regeneration respond to injury even in the absence of stem cells (*42–44*), we included cells from all three regeneration time courses in the calculation. On average, transcriptomes collected 0 – 4 dpa were ~45% of cells in each analyzed tissue sub-cluster (44.70% +/− 14.01%) (Fig. 4L). However, eleven tissue sub-clusters were disproportionately populated by cells from early timepoints – 72.72% of cells from 0 – 4dpa is 2 standard deviations above the average (dotted lines in Fig. 4D-L, Extended Data Fig. 8K). These sub-clusters were termed amputation-specific cell states and collectively included only 4.64% of cells in the atlas. Thus, these cell states were rare and may have been difficult to detect in bulk RNA-seq studies or single-cell reconstructions with fewer cells.

**Figure 4.**
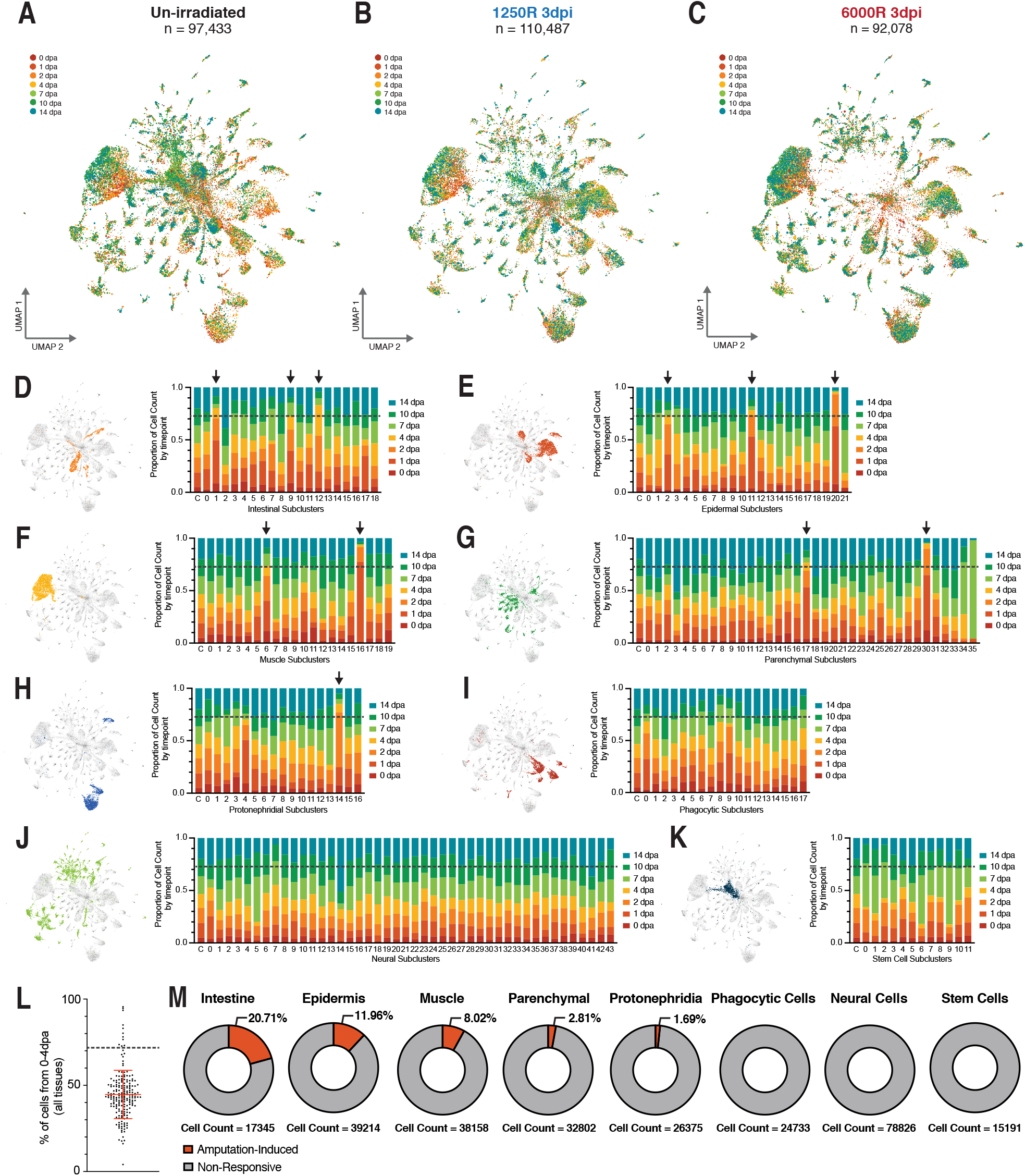
Amputation specific clusters are non-uniformly distributed across tissue lineages. UMAP embedding of captured single cell transcriptomes from biopsies taken from un-irradiated (A), sub-lethally irradiated (B), or lethally irradiated (C) animals, colored by time. Proportion of cells that fall into each timepoint within Intestinal (D), Epidermal (E), Muscle (F), Parenchymal (G), Protonephridial (H), Phagocytic (I), Neural (J), and Stem Cell (K) Sub-clusters. Black arrows indicate amputation-specific clusters. (L) % of cells in each cluster that come from 0dpa-4dpa. Error bars depict mean +/−SD. Dotted line indicates +2*SD. (M) Proportion of cell within each tissue that are in an amputation-specific cluster. An amputation specific cluster must have greater than 72.72% of sub-cluster cells captured from 0 dpa – 4 dpa (gray dashed line).

In order to determine which tissues were most responsive to amputation, we calculated the proportion of cells in each tissue belonging to an amputation-specific state. This analysis revealed that amputation-specific states were not uniformly distributed across tissues (Fig. 4M). Despite the need for regeneration of the anterior central nervous system, there were no neural transcriptional states that expanded during early time points after amputation. Instead, we found that a neural sub-cluster (neural cluster 5) expanded at day 7 and day 10 and was not produced in biopsies taken from irradiated animals (Fig4J, Extended Data Fig. 8B). This may indicate that the regeneration and remodeling of the central nervous system is a relatively late step during whole-body regeneration. Similar to the neural cell states, there were no amputation-specific cell states in the stem cell compartment, despite a well characterized activation of stem cell divisions after injury (*45, 46*). Finally, amputation-specific cells states were non-existent or minimally present amongst phagocytic, protonephridial (kidney-like), or parenchymal cells. Cells from amputation-specific states represented less than 5% of the total dataset, but ~8%, ~12%, and ~21% of cells captured from the muscle, epidermis, and intestine, respectively. Thus, amputation-specific transcriptional states were enriched in tissues representing all three germ layers – muscle (mesoderm), epidermis (ectoderm), and intestine (endoderm) (Fig. 4M). Given this enrichment, we focused on these tissues and their associated amputation-specific cell states for functional testing.

### Transient muscle cell states are required for tissue patterning after amputation

Muscle cells constitutively express regionally restricted position control genes (PCGs), which are required to maintain and regenerate the adult body plan (*47, 48*). As such, re-establishment of positional information in muscle cells is an important step required for regeneration of missing tissues in planaria (*42, 43, 49–51*). We identified two amputation-specific muscle cell states in our reconstruction of regeneration – muscle cluster 6 (MC6) and muscle cluster 16 (MC16) (Fig. 4F, Fig. 5B,C, black arrows). Notably, MC6 and MC16 represented only 0.785% (79/10,000) and 0.234% (23/10,000) of cells in the full dataset and only 8.02% of all muscle cells captured. Both MC6 and MC16 were upregulated 24 hours after amputation, but MC16 expansion was restricted to 24 – 48 hours after amputation and the expansion was less significant relative to homeostasis in biopsies taken from sub-lethally or lethally irradiated animals (Fig. 5D, Extended Data Fig. 9A-C, black arrows). We examined genes enriched in MC16 relative to other muscle clusters and found numerous genes known to regulate re-establishment of positional information after injury, including *follistatin* (*52, 53*), *notum* (*54*), *evi/wls*(*50*), and *glypican-1*(*42*). Genes enriched in MC16 were also largely restricted to the muscle and were wound-induced in all three single-cell regeneration time courses (Fig. 5E) and in a bulk RNAseq dataset (*55*) dataset of planarian regeneration (Extended Data Fig. 9D). An RNAi screen targeting MC16-enriched genes confirmed patterning phenotypes for *follistatin, notum, evi/wls*, and *glypican-1*. In addition, the screen identified *grp78* and *Ca-trans ATPase* as novel regulators of tissue homeostasis (Fig. 5G,H) and identified *junctophilin-1* as a novel regulator of tissue patterning (Fig. 5I,J).

**Figure 5.**
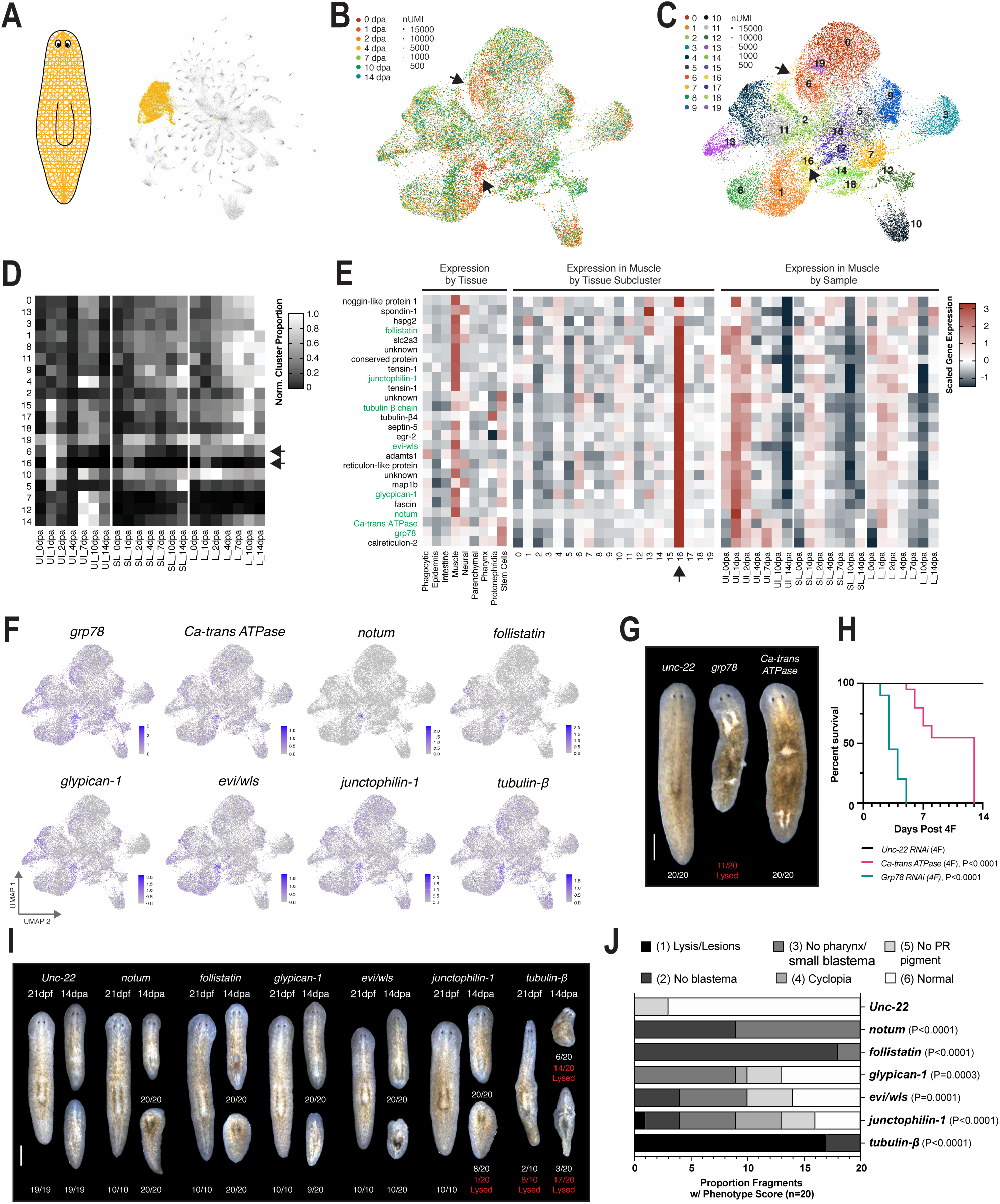
Regulators of tissue re-patterning were identified in wound-induced muscle clusters (black arrows). (A) UMAP embedding of global dataset with muscle cells highlighted. UMAP embedding of all muscle cells, colored by time after amputation (B) or tissue sub-cluster ID (C). (D) Scaled proportion of cells from each muscle sub-cluster across sampled conditions, normalized to sample in which sub-cluster had maximum representation. (E) Scaled mean expression of muscle cluster 16 enriched genes (black arrow) by tissue, muscle sub-cluster, and sample. (F) UMAP feature plots showing muscle expression of genes that produce penetrant RNAi phenotypes. (G) Representative images RNAi treated animals 3 days post feeding (H) Survival of RNAi treated animals shown in F (n=20 for each condition). (I) Representative images of homeostatic (21 days post feeding, n=10) and regeneration phenotypes (14 dpa, n=20) in RNAi treated animals. (J) Scoring of regeneration phenotypes. P values are Rank Log test (H) and Mann Whitney Test (J) compared to Unc-22 Control. Scale = 500um.

Our data support the existing model that muscle cells transiently express notum and other Wnt signaling regulators after injury to re-establish axial patterning. We also identified novel regulators of patterning and demonstrated that only a subset of muscle cells appear to be involved in re-establishing polarity signaling. Moreover, these activities occur even in the absence of stem cells or muscle progenitors, which is consistent with prior work showing that wound-induced expression of some PCGs occurs after lethal irradiation (*42, 43, 56*). Consequently, these data demonstrate that amputation induces a transient muscle cell state required for re-establishment of polarity signaling and subsequent patterning of missing tissue. This muscle cell state is rare and represents only a subset of muscle cells after injury.

### Injury-responsive Agat^+^ epithelial progenitors are required for tissue maintenance

In both vertebrate and invertebrate systems, the wounded epidermis expresses and secretes signaling molecules that promote wound closure and tissue regeneration (*10, 18, 19, 42*). In planaria, the molecules regulating differentiation of *piwi-1*^+^ progenitors into mature epidermal cells are some of the most well characterized amongst all tissue lineages. During normal tissue turnover, *zfp-1*^+^ sigma neoblasts progress through a *prog-2*^+^ early progenitor state and an *agat-3*^+^ late progenitor state. Cells then transition through a *zpuf-6*^+^ or *vim-1*^+^ stage before becoming mature *rootletin^+^, PRSS12*^+^, or *laminB*^+^ epidermal cells (*32, 57, 58*). Notably, *prog-2*^+^ and *agat-3*^+^ epidermal progenitors are some of the first cell types ablated after lethal irradiation, likely due to high levels of turnover in the epidermis (*57, 59*). We identified three amputation-specific cell states in the planarian epidermal lineage – epidermal cluster 2 (EC2), epidermal cluster 11 (EC11), and epidermal cluster 20 (EC20) (Fig.3E, Fig. 6A-C, black arrows). EC2, EC11, and EC20 contained 1.093% (109/10,000), 0.374% (38/10,000) and 0.097% (9/10,000) of cells in the full dataset, respectively, and represented only 11.96% of all epidermal cells captured. Genes enriched in amputation-specific epidermal states were specific to the epidermal lineage (Fig. 6D). The largest of these three amputation-specific clusters – EC2 – was characterized by high expression of *hadrian* (Fig. 6D), a wound-induced planarian gene expressed after injury in the mature epidermis (*42*). EC2 was induced transiently in biopsies taken from un-irradiated animals (only present at 24 and 48 hours post amputation (hpa)), but endured after 48 hpa in biopsies taken from sub-lethally or lethally irradiated animals (Extended Data Fig. 10A-D, black arrows). In contrast, EC11 and EC20 were induced at 24 and 48 hpa in biopsies taken from un-irradiated animals, but were significantly reduced in biopsies from sub-lethally irradiated animals, and not present at all in biopsies taken from lethally irradiated animals (Extended Data Fig. 10A-D, black arrows). Genes enriched in EC11 and EC20 included known markers of late epidermal progenitors, such as *agat-1, agat-2*, and *agat-3 (57, 59, 60*) (Fig. 6D, Extended Data Fig. 10F). We confirmed by in situ hybridization that *agat*^+^ cells are more densely packed adjacent to wound sites and that *agat-1, agat-2*, and *agat-3* are robustly expressed in these cells (Fig 6E, insets 1-3, Extended Data Fig. 10E).

**Figure 6.**
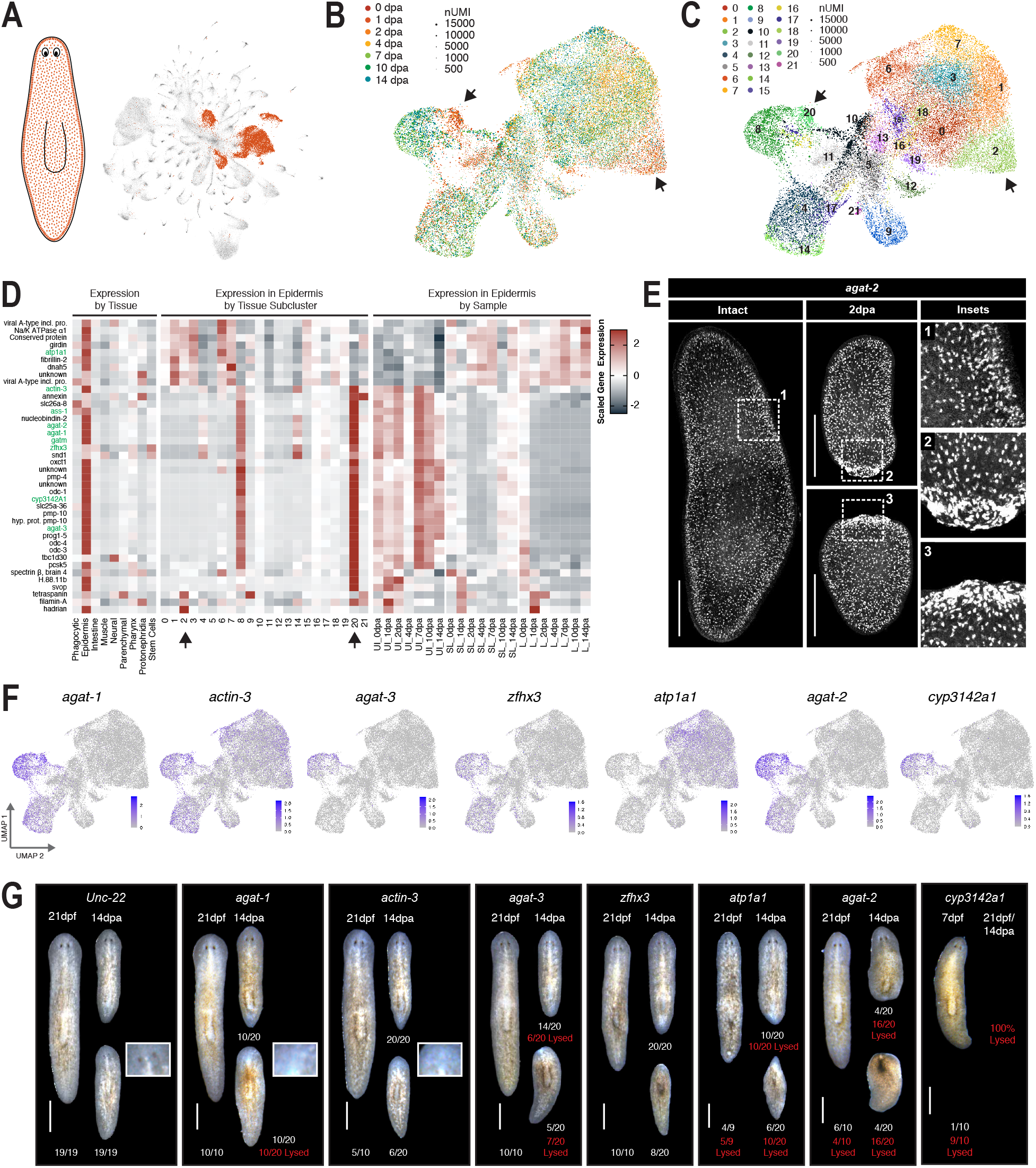
Amputation-specific states (black arrows) in the late epithelial progenitors are required for tissue homeostasis. (A) UMAP embedding of global dataset with epithelial lineage cells highlighted. UMAP embedding of all epithelial cells, colored by time after amputation (B) or tissue sub-cluster ID (C). (D) Scaled mean expression of genes enriched in wound-induced states (black arrows) by tissue, epithelial sub-cluster, and sample. (E) *agat-2* gene expression visualized by *in situ* hybridization (F) UMAP feature plots showing epidermal expression of genes that produced penetrant RNAi phenotypes. (G) Representative images of homeostatic (21 days post feeding) and regeneration phenotypes (14 dpa) in RNAi treated animals.

In order to test the function of amputation-specific epidermal cell states, genes enriched in EC2, EC11, and EC20 were depleted by RNAi and phenotypes were observed at homeostasis and after amputation. Surprisingly, a range of homeostasis and regeneration defects were observed in animals in which genes enriched in *agat-3*^+^ epidermal progenitors (EC20) were depleted (Fig. 6F,G, Extended Data Fig. 10G). *Agat-1* and *actin-3* RNAi-treated animals lacked photoreceptor pigmentation in posterior regenerates at 14 dpa (Fig 6G, insets). Animals treated with RNAi targeting *agat-3* or *zinc finger homeobox protein 3 (zfhx3*) had no blastema growth or reduced blastema growth at 14 dpa (Fig. 5F). Finally, three different genes produced lesions at homeostasis and no blastema or lesions after amputation – *cyp3142a1, agat-2*, and *sodium/potassium-transporting ATPase subunit α (atp1a1*) (Fig. 6G). While prior studies have identified regulators of epidermal differentiation expressed in *agat*^+^ cells, our results indicate that late epidermal progenitors also alter their transcriptional output in response to injury. Committed stem cell progeny have been shown to contribute to stem cell proliferation and tissue dynamics in other systems (*61*). Similarly, our results indicate that *agat*^+^ cells are not solely a progenitor state along a differentiation trajectory. Instead, this cell state is responsive to injury and functionally required for tissue maintenance and new tissue growth after amputation.

### Intestinal cells respond to injury and are required for regeneration

The endoderm is a well-established model for stem cell regulation during normal tissue turnover and in response to tissue injury (*15, 62, 63*). Several types of support and stem cells important for tissue maintenance and injury repair have been identified, including rare or transient cell states that only respond to particular types of injuries (*15, 17, 64, 65*). The planarian intestine contains three distinct cells types: secretory goblet cells, absorptive phagocytes/enterocytes, and recently identified ‘outer’ or ‘basal’ intestinal cells (*32, 66*). We found three different amputation-specific cell states within the planarian intestine – intestinal cluster 1 (IC1), 9 (IC9), and 12 (IC12) (Fig. 7A-C, Extended Data Fig. 11A-D). Consistent with amputation-specific cell states in other tissues, these cell states were very rare, representing 0.895% (90/10,000), 0.192% (19/10,000), and 0.110% (11/10,000) of cells in the full dataset. However, amputation induced cell states accounted for 20.71% of intestinal cells captured across the three regeneration time courses, indicating that the intestine is the most injury-responsive tissue in planaria. Expansion of IC1 and IC9 occurred at 24 hpa and persisted longer in biopsies taken from sub-lethally or lethally irradiated animals (Extended Data Fig. 11A-D, black arrows). Surprisingly, IC12 expanded slightly later after amputation (48-96 hpa) and was equally prevalent in biopsies taken from sub-lethally or lethally irradiated animals (Extended Data Fig. 11A-D, black arrows). The majority of genes enriched in IC1 were shared with more than three other intestinal clusters and these genes were particularly shared with IC2 and IC9, so we focused on IC9 and IC12 for functional testing. Significantly, IC9 and IC12 markers revealed the cell states to be similar to previously described ‘outer’ or ‘basal’ intestinal cells and absorptive enterocytes, respectively (Fig. 7D, E, Extended Data Fig. 11E, se also (*32, 66*)). Genes enriched in amputation-specific intestinal clusters included methyltransferases (*pbrm-1, mettrans*), regulators of endocytosis (*myosin 1E*), regulators of lysosomal degradation (*cathepsin B, cathepsin L*), and many regulators of gut function and metabolism (*lectin2b, lipase, TDO, PEPCK, glycoside hydrolase*).

**Figure 7.**
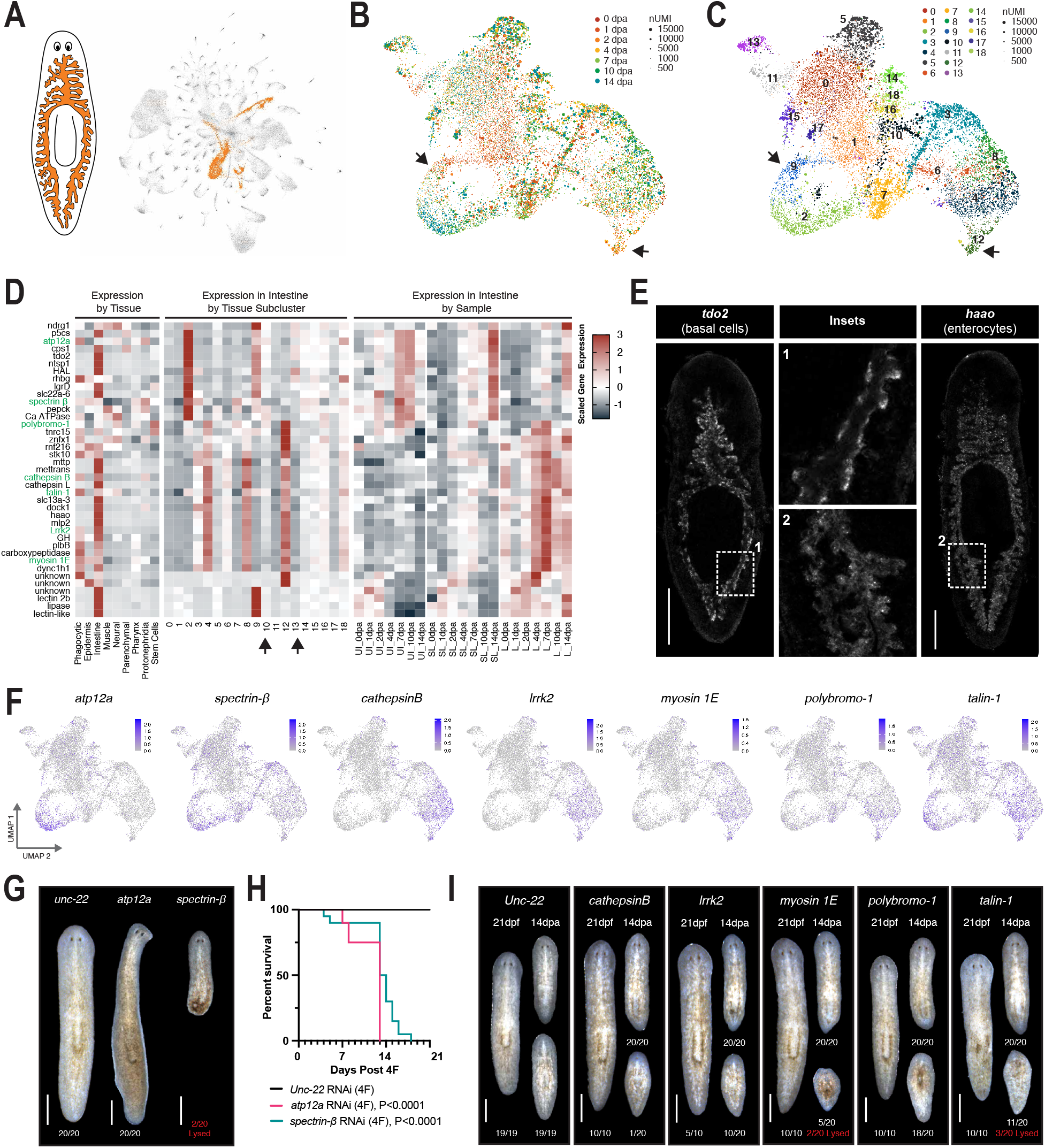
Amputation-specific states (black arrows) in the intestine are required for regeneration. (A) UMAP embedding of global dataset with intestinal cells highlighted. UMAP embedding of all intestinal cells, colored by time after amputation (B) or tissue sub-cluster ID (C). (D) Scaled mean expression of genes enriched in wound-induced states (black arrows) by tissue, epithelial subcluster, and sample. (E) *tdo2* and *haao* gene expression visualized by *in situ* hybridization (F) UMAP feature plots showing intestinal expression of genes that produced penetrant RNAi phenotypes. (G) Representative images RNAi treated animals 3 days post feeding. (H) Survival of RNAi treated animals shown in F (n=20 for each condition). (I) Representative images of homeostatic (21 days post feeding, n=10) and regeneration phenotypes (14 dpa, n=20) in RNAi treated animals.

In order to test whether amputation-specific intestinal cell states were required for planarian regeneration, genes enriched in IC9 and IC12 were depleted using RNAi. Several defects in tissue homeostasis and regeneration were observed (Fig. 7F-I, Extended Data Fig. 11G). Notably, RNAi depletion of *atp12a* and *spectrin β* (both enriched in IC2/9 – ‘outer/basal’ cells) resulted in lesions or inch-worming at 3 days post feeding (Fig. 7G) and lysis before 21 days post feeding (Fig. 7H). In contrast, RNAi depletion of *myosin1E, pbrm-1*, or *talin-1* (all enriched in IC12 – enterocytes) resulted in small blastemas and limited or no pharynx regeneration (Fig. 7I, Extended Data Fig. 11G). RNAi depletion of *cathepsin B* and *lrrk2* (also enriched in IC12) resulted in an increased incidence of cyclopia or un-pigmented photoreceptors (Fig. 7I, Extended Data Fig. 11G). Notably, tail regeneration in anterior fragments was less affected in all RNAi treatments. These data support a model in which amputation induces changes in transcriptional output for most ‘outer/basal’ intestinal cells and a rare subset of absorptive enterocytes (Fig. 7B,C, black arrows). This may reflect changes in metabolism required for whole-body regeneration. In addition, we observed that ‘outer/basal’ intestinal cells are more sensitive to irradiation than other intestinal cell types, perhaps due to higher rates of turnover within the intestine or high signaling feedback with nearby stem cells. This is supported by the observation that RNAi of genes enriched in IC2 and IC9 produced homeostatic lesions, while RNAi of IC12-enriched genes produced less severe regeneration defects.

Over 20% of cells captured from the intestine occupied amputation-specific cell states. Nearly all ‘outer/basal’ cells shifted their transcriptional output to modify metabolic signaling and upregulate lectin-like molecules after injury and a small subset of enterocytes altered gene expression, perhaps via increased expression of the chromatin modifier *pbrm-1*. Depletion of genes enriched in amputation-specific intestinal states produced homeostatic and regeneration defects. We conclude from these results that the intestine is the most transcriptionally responsive tissue in planaria during whole-body regeneration and that amputation-specific signaling in a subset of intestinal cells is required for tissue maintenance and new tissue growth in planaria.

## Discussion

To date, the study of regenerative capacity has largely focused on signaling mechanisms near the injured tissue required for stem cell proliferation and differentiation following injury (*15, 19, 67–70*). Relatively little is known about how post-mitotic cells contribute to whole-body regeneration, particularly from cell types that may act on stem cells and their immediate progeny from a greater distance (*17, 18, 71*). Here, we present a comprehensive single cell reconstruction of whole-body regeneration over time. To ensure the dataset contained only cellular components essential for regeneration, minimal body fragments were taken from unirradiated, sub-lethally irradiated, and lethally irradiated planarians. This strategy allowed us to define the cellular dynamics of successful and un-successful regeneration across an entire animal for the first time and to identify rare amputation-specific cell states. Additionally, we determined which cellular states were or were not dependent upon stem cell proliferation and functionally tested which were required for complete replacement of all missing tissues.

Amputation-specific cell states identified in the mesoderm (*notum*^+^ MC16), ectoderm (agat3^+^ EC20), and endoderm (*tdo*^+^ IC9 and *pbrm-1*^+^ IC12) were required for regenerative capacity (Figs. 5-7). These cell states were both rare and transiently induced only after injury (Fig. 4). In the muscle, only a subset of cells after amputation induced the patterning cues (*notum, follistatin, evi/wls, glypican-1, junctophilin-1*) required for re-establishment of the body plan and subsequent growth of missing tissues (Fig. 5). In the epidermis, we discovered a previously unknown *agat-3*^+^ cell state induced by injury and required for tissue homeostasis (Fig. 6). We also found that ‘basal/outer’ intestinal cells and a rare subset of enterocytes in the intestine (Fig. 7C, black arrows) undergo a transient change in transcriptional state required for tissue maintenance and new tissue growth, respectively (Fig. 7G-I). Importantly, wound-induced muscle (MC16) and wound-induced enterocytes (IC12) were generated in tissue fragments independent of stem cell proliferation (Extended Data Fig. 9A-C, Extended Data Fig 11A-C), consistent with previous observations that a large percentage of wound-induced genes do not require new proliferation (*42, 44*). In all cases, amputation-specific cell states required for regeneration amounted to less than 1% of captured cells and were only present at a few time points after amputation (Fig. 2, 4). In other systems, an increase in transcriptional plasticity or activation of rare cell types can be required for wound healing, particularly in systems where stem cells have been ablated (*64*). Our results reveal that regenerative capacity may arise from an organism’s ability to produce these rare and transient cell states within a subset of differentiated tissues after injury. We refer to these as transient regeneration-activating cell states (TRACS).

The observation that TRACS are both transient and rare may have important implications for the regulation of growth capacity. TRACS were only present at specific times after amputation and were transcriptionally distinct from cell states present at homeostasis. This transient activation may act as a guard against un-checked growth in the absence of injury. Indeed, some genes expressed in amputation-specific intestinal TRACS (*pbrm-1* and *TDO*) have already been implicated in cancer proliferation and tumor immune evasion (*72–76*). Therefore, it is possible that mechanisms ensuring the transiency of regeneration-specific cell states and mechanisms that prevent oncogenesis in aging tissues may share underlying principles. TRACS also occurred in rare subsets of differentiated tissues. The rarity of these cell states may account for why regenerative capacity is so unevenly distributed across the animal kingdom. A rare cell state is more easily lost during evolution and – perhaps – more easily reactivated (*77*). Several planarian species incapable of head regeneration have been described and regenerative capacity within these species restored by inhibition of Wnt signaling (*78, 79*). We can now hypothesize that this reduction in Wnt signaling may have restored the ability of rare, amputation-specific muscle TRACS (*notum*^+^ MC16) to re-program the planarian body plan. If generally applicable, re-activation or transient restoration of regeneration-specific cell states could unleash regenerative capacity in systems where such capacities are presently limited.

The results presented in this study characterize the cellular components required for regeneration at unprecedented molecular resolution, facilitating the discovery of rare cell states (TRACS) and novel molecules required for regeneration. Moreover, these data support a model in which regenerative capacity in general may be linked to the activation of transcriptional plasticity in rare differentiated cells.

## Methods

### Animal Husbandry

*S. mediterranea* animals from the asexual clonal strain CIW-4 (C4) (*79*) were maintained in 1X Montjuic salts in a planarian re-circulation culture system as previously described (*80*). Tissue biopsies were taken from large animals fed chunk beef liver three times per week in a recirculation system and starved for 3 – 4 days in static culture prior to amputation. Small animals utilized for *in situ* hybridizations and the RNAi screens were fed chunk liver once a week in a recirculation system and starved in static culture at least 4 weeks prior to experimental use. For irradiation treatments, worms were dosed with gamma irradiation in an MDS Nordion Gammacell 40 Exactor low dose-rate research irradiator. Delivered dose was calculated by internal software correcting for decay, and an independent second timer verified exposure time. Dose rate and distribution are professionally monitored annually.

### Tissue biopsies

For tissue biopsies, animals between 1 and 2cm in length and wider than 2mm were picked from recirculation culture systems and starved 3 – 4 days in static culture. Animals were anesthetized using cold chloretone solution (0.1-0.2% w/v chloretone in 1X Montjuic salts) for 4 – 5 minutes or until immobile. Animals were rinsed with cold Holtfreter’s solution (3.5g/L NaCl, 0.2g/L NaHCO3, 0.05g/L KCl, 0.2g/L MgSO4, 0.1g/L CaCl2, pH 7.0-7.5) and placed on Whatman #3 filter paper saturated with cold Holtfreter’s to minimize movement (*81*). Limited reuse biopsy punches (World Precision Instruments – #504528 [0.5mm], #504529 [0.75mm], #504646 [1.0mm], #504530 [1.2mm], #504647 [1.5mm]) were used to extract round tissue fragments from the center of the animal’s posterior tail tissue and the biopsy plunger was used to deposit the fragments into cold Holtfreter’s solution to wound heal. Tissue fragments were stored at 20°C overnight and Holtefreter’s solution was exchanged for 1X Montjuic + 50μg/mL gentamycin at 24 hours post amputation.

### Generation of fixed planarian single-cell suspensions

To generate a cell suspension, animals or biopsies were rinsed in calcium-magnesium free buffer with 1% BSA (CMFB) (*82*), finely chopped with a single edge stainless steel blade, then agitated on a benchtop rocker in CMFB for 20 – 30 minutes with vigorous pipetting every 3-5 minutes. After maceration, cells were filtered through a 30μm Filcon cell-strainer and centrifuged for 5 minutes at 300g. Cells were re-suspended in 1X CMFB + 1:1000 Live Dead Dye (Life Technologies L34975) and incubated on ice for 20 minutes. Cells were centrifuged for 5 minutes at 300g and resuspended in 1mL PBS + RNase Inhibitors (Invitrogen AM2694, Qiagen Beverly Inc Y9240L) and fixed as previously described for split-pool ligation-based RNAseq. Briefly, 3mL 1.33% paraformaldehyde in PBS was added to 1mL of suspended cells and suspension was incubated on ice for 10 minutes. Then 160μL of 5% Triton-X was added to solution, gently pipetted, and cells were incubated on ice for 3 minutes. Cells were then centrifuged at 1000g for 10 minutes, resuspended in 500μL PBS + RI, and then mixed gently with 500μL cold 100mM Tris-HCl, pH 8.0. Cells were centrifuged for 10 minutes at 1000g, re-suspended in 1mL 5μg/mL Hoescht 33342 in 0.5X PBS + RI, incubated at RT for 10 minutes, and stored on ice prior to FACS.

### Image Cytometry

Image cytometry was used to determine the penetrance and localization of SPLiT-seq reagents in dissociated and fixed planarian cells. Linker oligonucleotides tagged with Atto dyes were purchased from IDT to visualize round 2 and round 3 barcoding reagents. In addition, biotin tagged barcoded molecules were visualized using a FiTC-tagged streptavidin (Extended Data Fig. 2A). At each successive step of oligonucleotide barcoding, samples were analyzed by imaging flow cytometry to assess by signal intensity of labeled oligonucleotides whether reagent penetration had occurred in dissociated intact cells and/or in cellular debris.

Data was acquired on an ImagestreamX mkII instrument at 60 running at the low/sensitive flow rate. Channels 1, 2, 7, 10, 11 and 12 were used to collect signal from BF, FITC, Hoechst, Atto 565, Atto 633, and live/dead780 respectively. Bandpass filters for probes were 528/55 (Ch2), 457/45 (Ch7), 610/30 (Ch 10), 702/85 (Ch 11), and 762/35 (Ch 12). Following imaging, spectral compensation was performed using files collected from single-stained samples for each fluorophore and spillover coefficients were calculated in IDEAS software v6.2.189. Analysis for the presence of probe intensity above background levels was also performed in IDEAS by setting gating regions, first on Hoechst positive and brightfield area dot plots to filter out non-cellular events lacking a DNA signal, then by setting gating regions on negative control samples (no probe or no secondary stain) and reporting the frequency of events above the level of negative control samples.

### Methods for split-pool ligation-based RNAseq (SPLiT-seq) of fixed planarian cells

Dissociated planarian cells stained with the Near-IR Live/Dead Dye and Hoescht 33342 were sorted with an influx sorter using a 100μm tip. Hoechst was excited with 100mW of 355nm UV and collected behind a 450/20 bandpass. Near-IR Live/Dead dye was excited with 631nm and collected behind a 750LP filter. Sorting was done in purity mode at approximately 12,000-15,000 eps. For each sample, 400,000 – 500,000 single Hoescht+/DeadDye-cells were collected for barcoding. Split-pool ligation-based barcoding of dissociated cells was performed as previously described (*31*), see also https://sites.google.com/uw.edu/splitseq/protocol.

In brief, cells from each sample were distributed into 48 wells containing reverse transcription reaction buffer and barcoded oligo(dT) and hexamer primers. The first barcode was added via the completed reverse transcription reaction. Cells were pooled together, centrifuged, resuspended in NEB 3.1 Buffer, and then distributed into a 96-well plate containing a T4 ligase ligation mix and DNA oligo barcodes. The ligation reaction was incubated for 30 minutes at 37°C with gentle rotation, then for another 30 minutes at 37°C following the addition of a solution containing blocking oligo strands. Cells were collected from the plate, gently mixed, and distributed into a second 96-well plate, again containing DNA oligo barcodes. The ligation of the third barcode was achieved during another 30 minute incubation at 37°C with gentle rotation, followed by the addition of a blocking solution, again containing a blocking DNA oligo to bind to un-ligated DNA oligo barcodes. All cells were pooled and the reaction terminated with EDTA. Cells were centrifuged at 1000g for 10 minutes in a swinging bucket centrifuge and resuspended in 1X PBS + SUPERase In RNase Inhibitor (Invitrogen AM2694). Single, intact Hoescht+ cells were sorted with an influx sorter using a 100μm tip and 150,000 – 200,000 cells were re-suspended in 400μL 1X PBS + SUPERase In RNase Inhibitor. Cells were split into 8 sub-libraries, with 50μL of suspension going into each tube. 50μL lysis buffer (20mM Tris, pH 8.0, 400mM NaCl, 100mM EDTA, pH 8.0, 4.4% SDS) and 10μL proteinase K (20mg/mL) was added to the barcoded cells. Lysates were incubated at 55°C for 2 hours with 200rpm agitation, then stored at 70°C until library preparation.

### Library preparation and sequencing

Having been frozen at −70°C, cell lysates were thawed and sequencing libraries generated according to the SPLiT-seq Protocol with some modifications. (v3.0) (*31*), see also https://sites.google.com/uw.edu/splitseq/protocol) Incubation plus agitation steps were performed at room temperature with 500 RPM agitation on a thermomixer (with the final, 42°C, template switching incubation agitated at 300 RPM). Following template switch clean-up, all steps were conducted in 96-well plate format. Monitored cDNA amplification was stopped once signal left exponential phase (10 cycles), and a SPRI bead clean-up performed using two additional (200μL) ethanol washes. Amplified cDNA was checked for quality and quantity using an Agilent 2100 Bioanalyzer and Invitrogen Qubit Fluorometer and normalized to 600pg per sample for sequencing library preparation using Illumina Nextera XT library preparation kit. The PCR amplification program was modified to incorporate an initial 3 min 72°C hold for increased yield. Resulting short fragment libraries were assessed for quality and quantity, pooled equal molar in batches of eight, and sequenced on a total of nine High-Output flow cells (three for each regeneration time course) of an Illumina Next Seq 500 instrument using NextSeq Control Software 2.2.0.4 with the following paired read lengths: 66 bp Read1, 6 bp I7 Index and 94 bp Read2. Following sequencing, Illumina Primary Analysis version NextSeq RTA 2.4.11 and bcl2fastq2 v2.20 were run to demultiplex reads for all libraries and generate FASTQ files.

### Data processing and alignment of SPLiTseq dataset and public Drop-seq dataset

Pooled splitseq libraries were demultiplexed into individual timepoint samples using custom python script (Available on github upon publication). Read2 contains three rounds of cell barcodes which were designed during library prep and pooling step (UMI: 1-10bp, cell barcode1: 87-94bp, cell barcode2: 49-56bp, cell barcode3: 11-18bp). All samples were demultiplexed allowing up to one mismatch per cell barcode round, with quality score more than 10. Each timepoints samples were then processed using Drop-seq tools software (http://mccarrolllab.org/dropseq/) for barcode extraction, gene annotation and cell expression matrix generation. All samples were aligned using STAR aligner with the following parameters (--outFilterMatchNmin 0 --outFilterMismatchNmax 10 --outFilterMismatchNoverLmax 0.3 -- outFilterMismatchNoverReadLmax 1). Uniquely aligned reads were then annotated using Drop-seq tools function TagReadWithGeneExon based on planarian gene interval refFlat file. Final UMI expression matrix was collapsed using Drop-seq tools function DigitalExpression with parameter “EDIT_DISTANCE=1”. Raw and processed data files can be retrieved from the GEO database under accession number SE146685. Public Drop-seq reference datasets were downloaded via GEO accession GSE111764. Data were processed using the same functions and parameters with SPLiTseq datasets as described above.

### Quality filtering, data normalization, and clustering of cells

Low quality cells from the raw sequencing (UMI < 500 and/or Genes < 75) were excluded from subsequent analysis. The average numbers of gene and UMIs from each sample following this quality filtering are listed in Supplementary Figure 3C. Utilizing R’s Seurat 3 Package (*83*) version 3.1.1, we used SCTransform (*84*) with the default settings to normalize and variance stabilize the data. The 3000 most highly variable genes were used for principal component analysis of the SCTransformed data. 150 principal components (PCs) were calculated using Seurat’s RunPCA function. RunUMAP was then used to generate both two-dimensional and three-dimensional UMAP embeddings (*85*). FindNeighbors was run using 150 PCs to generate a shared nearest neighbor graph and FindClusters (all default settings) was used to cluster the data, resulting in 89 clusters designated “0” through “88.”

### Identifying enriched transcripts and tissue annotation

We used Seurat’s FindAllMarkers with default parameters to find marker transcripts for each of the 89 global clusters, resulting in a total of 3,558 markers comprised of 1,015 unique transcripts (some transcripts were markers for more than one cluster). Gene annotations were sourced from the Sánchez Alvarado lab’s Rosetta Stone Transcript Mapper (*86*). The most significant 20-40 transcripts enriched in each global cluster were mapped to dd_Smed-v6 or dd_Smed_v4 using the same mapper and their expression and tissue specificity was characterized in published planarian single-cell datasets (*32, 33*). Most global clusters were easily assigned to a previously described tissue lineage due to a shared gene expression profile and in many cases, a cluster could be matched 1-to-1 to a cluster or sub-cluster in a previously published dataset.

In addition to manually annotating each global cluster, we used Seurat’s TransferData function (*82*) to classify cells in our dataset by tissue using one of the prior planarian single-cell studies (*32*) as a reference for tissue annotation. Raw data (GSE111764) was aligned using STAR aligner using the same parameters outlined above and a gene expression matrix generated, again using Drop-seq_tools-1.13. Only cells which could be unambiguously identified in the published Fincher metadata using a combination of cell barcode and the type of tissue sample were retained. Tissue annotations from the published metadata (*32*) were added. FindIntegrationAnchors (dims=1:30) was used to identify “anchor” cells, then IntegrateData (dims=1:30) was used to integrate the published dataset with our full SPLiT-seq dataset. PCA and UMAP embedding were performed with 30 PCs. FindTransfer Anchors (dims=1:30) found transfer anchors using the published data as the reference and our dataset as the query. TransferDataset used the identified transfer anchors and the published metadata’s tissue annotations to predict tissue annotations for all cells in our SPLiT-seq dataset. The predicted tissue type for each cell was then added to the SPLiT-seq metadata.

Global cluster 1 contained very few significantly enriched genes and most enriched genes were also highly expressed in cells from the epidermis, muscle, or phagocytic cell lineages. In prior work, clusters lacking specific markers were labeled an artifact and excluded from subsequent analysis (*32*). However, we found that cells in cluster 1 were not uniformly distributed across all samples. Instead, they were more common in early time points in all three regeneration time courses, suggesting that the transcriptional state might be a wound-induced state characterized by a lack of markers. We therefore included cluster 1 in the full dataset and labeled it ‘Non-differentiated.’ Cells from this cluster were not assigned to any tissue subset or included in analysis of cells/sub-clusters from known tissue lineages.

### Sub-clustering of tissue subsets

Tissue annotations were added to the metadata for all cells in the dataset, after which the data was broken into subsets by tissue type. For each resulting tissue subset, the raw RNA counts were normalized with SCTransform as before prior to performing PCA with the number of PCs determined on a tissue-by-tissue basis. The global clusters combined into each tissue subset and the PC parameters for tissue level sub-clustering are summarized in the following table.

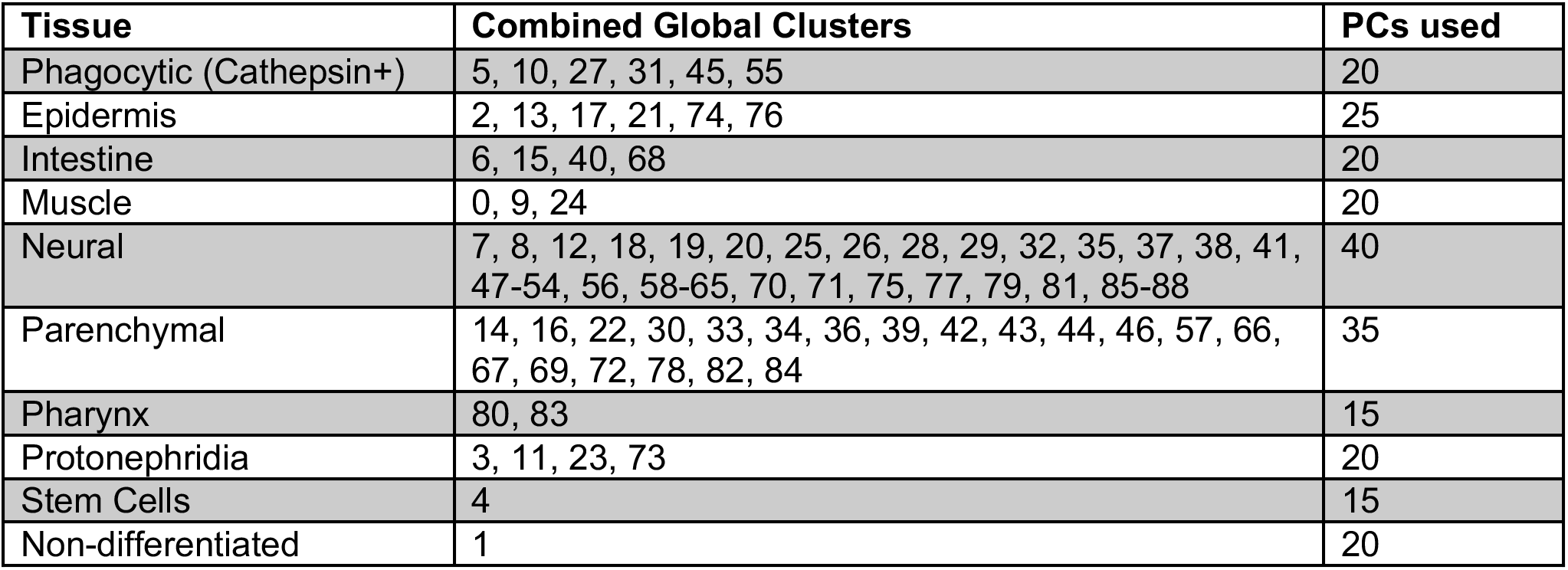

As before, we generated both two- and three-dimensional UMAP embeddings, a shared nearest neighbor graph using the same number of PCs as in the principal component analysis, and clustered using the default settings. We then used FindAllMarkers with default settings to find markers for each of the tissue-level clusters (“sub-clusters”). This resulted in a total of 211 sub-clusters (18 Phagocytic/Cathepsin+, 22 Epidermis, 19 Intestine, 20 Muscle, 44 Neural, 36 Parenchymal, 8 Pharyngeal, 17 Protonephridia, 12 Stem Cell, and 15 Non-differentiated).

### Proportional down-sampling for global data visualization

Because the number of cells in each sample varied from 3,461 to 36,344, UMAP visualizations became dominated by the larger samples, affecting the relative abundance of cells in each cluster. While the relative abundance of cells in each cluster could be normalized by sample quite easily in tissue proportion heat plots, this was more difficult in UMAP plots. To mitigate this difficulty, we created a proportionately down-sampled dataset with a target maximum of 6000 cells per sample. Both the un-irradiated and sub-lethally irradiated day 0 samples had fewer than 6000 cells, so the entire sample was included. For each of the other samples, we calculated the relative proportion of each global cluster and multiplied by 6000 to determine the number of cells to retain from each cluster. The desired number of cells was then randomly selected from among the cells in that cluster in the given sample. When needed, we rounded all cell counts up to the next equal or greater integer, resulting in more than 6,000 cells per sample in the final resulting sub-sampled dataset. This sub-sampled dataset was used as the reference object for most global UMAP plots (Fig 2A-C, Tissue highlight plots, Extended Data Fig. 3B). All other plots (cluster proportion, scaled gene expression, etc.) were produced from calculations on the full dataset containing all 299,998 cells.

### Cloning of *S. mediterranea* transcripts

The Sánchez Alvarado lab consensus transcriptome was used as the reference sequence for designing PCR primers to amplify transcripts of interest. Primer3 was used to design PCR primers to amplify target regions 400-500 nucleotides in length from cDNA template. Overhangs homologous to the pPR-T4P vector were added to 5’ ends of each primer. PCR products were inserted into linearized vector using Gibson Assembly. Reactions were transformed directly into *Eschericia coli* strain HT115. Insert sequences were verified by sequencing.

### In situ hybridizations

For RNA expression analysis in whole planarians, NBT/BCIP and Fluorescent in situ hybridizations were performed as previously described (*87, 88*). Following hybridization, DIG-probes were detected with antibodies in MABT containing 5% horse serum appropriate for NBT/BCIP in situ hybridization (Roche anti-DIG-AP 1:1000) or for FISH (Roche anti-DIG-POD 1:1000). NBT/BCIP developed animals were stored in 80% glycerol overnight prior to mounting, while FISH animals were soaked overnight in a modified ScaleA2 solution (40% glycerol, 2.5% DABCO, 4M urea) prior to mounting and imaging.

### RNAi food preparation and feeding

Cloned gene vectors transformed into *E. coli* strain HT115 were cultured in 24-well, round-bottom culture plates in 2X YT bacterial growth media with 50μg/mL Kanamycin and 10μg/mL Tetracycline for 16 – 18 hours at 37°C. Production of dsRNA was induced by adding 6mL 2X YT bacterial growth media with 50μg/mL Kanamycin, 10μg/mL Tetracycline, and 1mM IPTG. Bacteria were cultured for an additional 4 hours at 37°C following induction. Beef liver was homogenized by adding 800μL 0.2X food coloring in Montjuic Salts to 2g beef liver puree, followed by pipetting until homogeneous. Cultures were centrifuged 10 minutes at 1500g, supernatant removed, and bacterial pellets were resuspended in 60μL homogenized beef liver. Planarians were fed RNAi food four times with two days between each feeding (1.5uμL RNAi food per 3 – 5mm planarian worm). Water exchanges were performed after each feeding and dish exchanges along with full water exchanges performed 24 hours prior to each new feeding. For all RNAi screens, *unc-22* dsRNA was used as a negative control and *follistatin* dsRNA was used as a positive control for failed regeneration.

### Phenotyping after RNAi

Following RNAi feedings, animals were moved to clean dishes with fresh Montjuic solution + 50μg/mL gentamycin. In the primary screen, animals were imaged 7 days post feeding for homeostatic phenotypes, then all surviving animals were bisected through the pharyngeal region to produce anterior and posterior fragments. Regeneration phenotypes were visually inspected and scored 7 days post amputation and animals were imaged and scored 14 days post amputation. For the secondary screen, the number of animals was doubled (n=20). RNAi depletions that had produced lesions by 3 days post feeding in the primary screen were assigned to survival curve assays, while phenotypes that had produced regeneration phenotypes or homeostatic defects only after amputation were assigned to regeneration assays. For survival curves, animals were monitored daily after the fourth feeding and lysis events noted until all animals were dead. A Log-rank (Mantel-Cox) test was used to compare survival of all RNAi conditions to the Unc-22 RNAi control. For regeneration assays, animals were imaged 7 days post feeding. 10 animals were bisected to produce anterior and posterior fragments, while 10 were left intact. At 14 days post amputation or 21 days post feeding, homeostatic and regeneration phenotypes were imaged and scored. Mann-Whitney U tests were used to compare RNAi conditions to Unc-22 RNAi controls. Results of primary and secondary screens are provided in Supplementary Table 3.

### Microscopy

Colorimetric whole-mount in situ hybridization samples and live worm or fragment images were acquired using a Leica M205 microscope. Following image acquisition, non-tissue background was subtracted, contract and image intensity adjusted, and the edited image was converted to grayscale for data presentation. No quantifications were performed on contrast or intensity adjusted images and all raw, original data is available in the original data repository (http://www.stowers.org/research/publications/libpb-1513). Confocal images of fluorescent in situ hybridization samples were acquired using an LSM-700 inverted confocal microscope. Stitching of tiles for whole worm images was performed with Fiji plugins (grid/collection stitching) using custom macros for batch processing. Maximum intensity projections of the stitched z-stacks were generated to visualize gene expression across the entire animal, then rotated, cropped and inverted for data presentation (Extended Data Fig. 12G and Extended Data Fig. 13G).

## Data and materials availability

Original data underlying this manuscript can be accessed from the Stowers Original Data Repository at http://www.stowers.org/research/publications/libpb-1513. SMEDIDs for all cloned genes are listed in Supplementary Table 3 and sequence information can be found at https://planosphere.stowers.org/find/genes. All single cell RNAseq data and cell by gene matrices used to generate the figures in this manuscript have been deposited in the Gene Expression Omnibus Database under the accession code GSE146685.

## Code Availability

Original scripts used for the analysis and visualization of single-cell sequencing data is available at https://github.com/0x644BE25/smedSPLiT-seq.

## Acknowledgments

We thank members of the A.S.A. laboratory for discussion and advice. We are grateful to the Stowers cytometry and molecular biology core facilities for technical contributions and methods development.

## Author contributions

Conceptualization and data interpretation (BWBP, ASA), data analysis (BWBP, CB, SC), acquisition of data (BWBP, AMK, AS, AB), cloning of planarian gene transcripts (FGM), writing of original manuscript (BWBP), supervision and funding acquisition (ASA), and revision and editing of manuscript (all authors).

## Materials and Correspondence

Correspondence should be directed to either.

## Supplemental Figures

**Extended Data Fig. 1.**
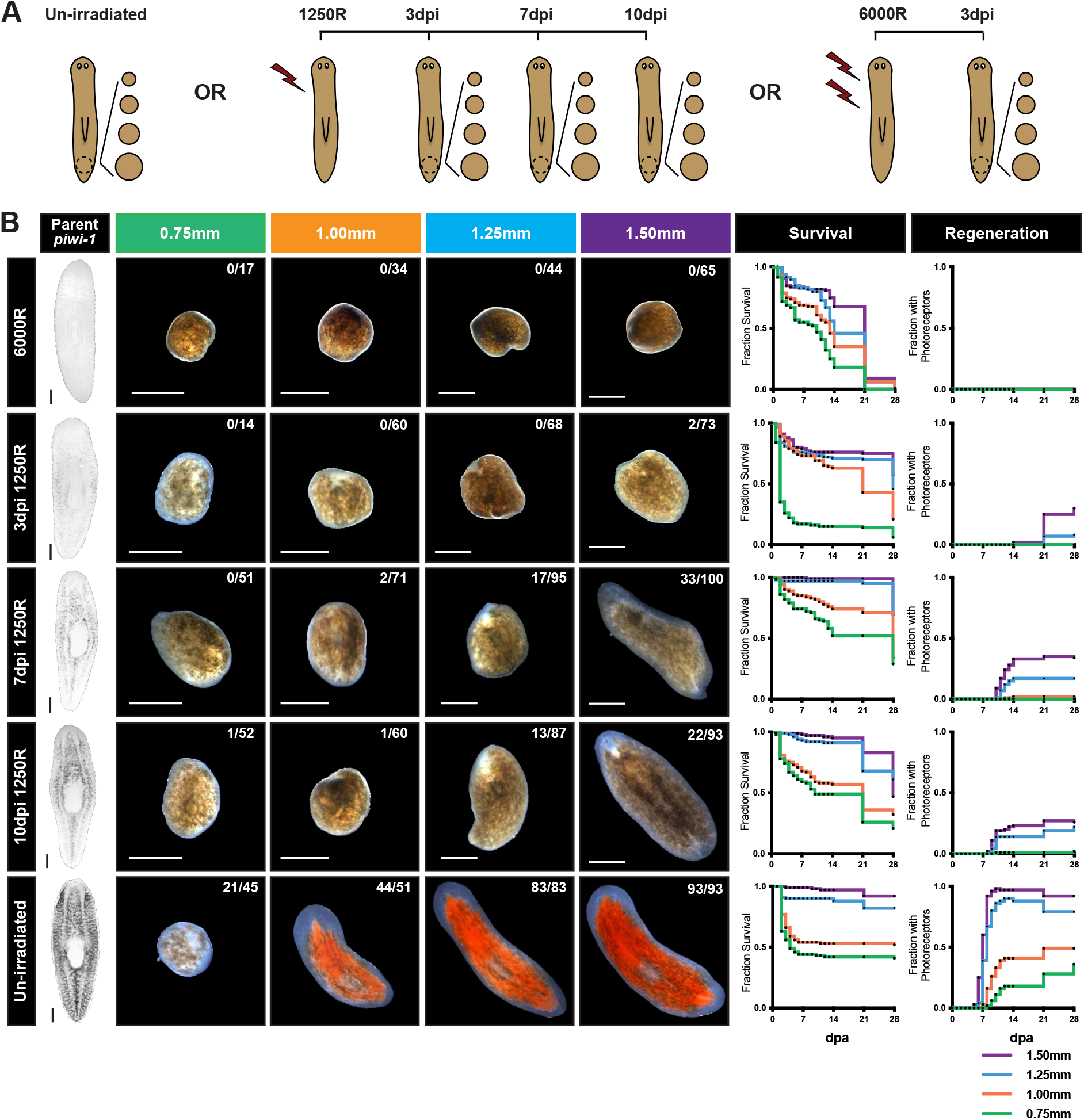
Survival and regeneration of 0.75mm – 1.50mm biopsies taken from unirradiated, sub-lethally irradiated, and lethally irradiated animals. (A) Schematic of experimental design. (B) Piwi-1 expression of parent animals at time biopsy was taken after irradiation treatment (right), as well as representative images 14 days post amputation (dpa), survival curves, and scoring of regeneration of photoreceptor pigmentation of biopsies 0.75mm – 1.50mm taken from comparable parent animals following irradiation treatment. Notation on representative images indicates number of fragments that regenerated photoreceptors by 14dpa out of total surviving at 14dpa (n=96-108 biopsies). Scale = 500um.

**Extended Data Fig. 2.**
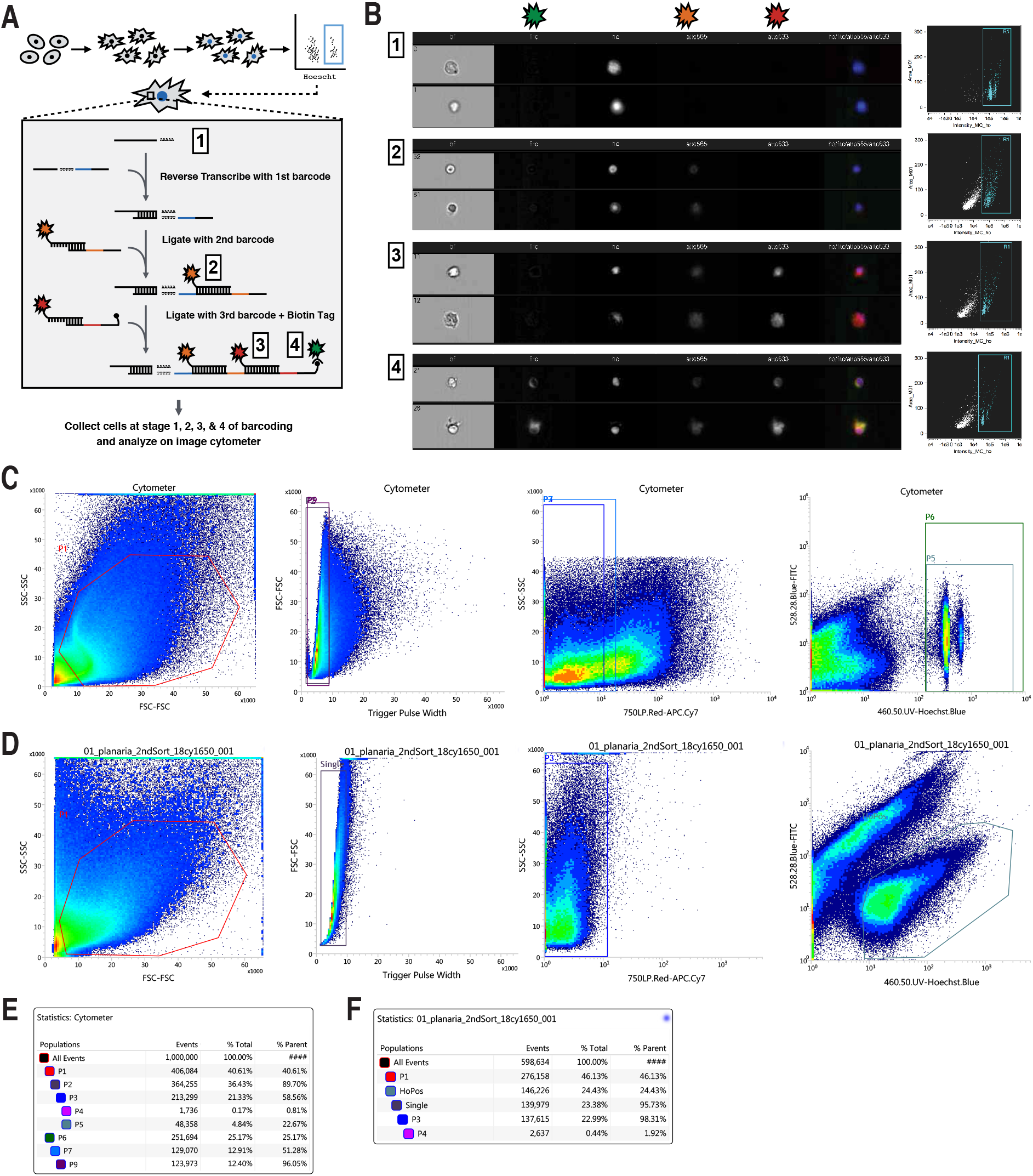
(A) Schematic of experimental design using Atto-conjugated linker molecules to visualize SPLiTseq reagents after second [2] and third [3] round barcoding, and to detect biotin tagged molecules [4]. (B) Representative images of cells/objects detected in the Hoescht+ compartment at all steps of the barcoding process and Area vs. Hoescht Intensity plots with Hoescht+ cell compartment highlighted from each stage of barcoding. Note the accumulation of non-nucleated debris that occurs during the barcoding process that needed to be removed prior to sequencing. As a result, Hoescht+ intact cells were sorted following barcoding using the plot depicted in B [4] as a guide. Gating strategy utilized pre-barcoding (C) and post-barcoding (D). Abundance of sorted population pre-barcoding (E) and post-barcoding (F).

**Extended Data Fig. 3.**
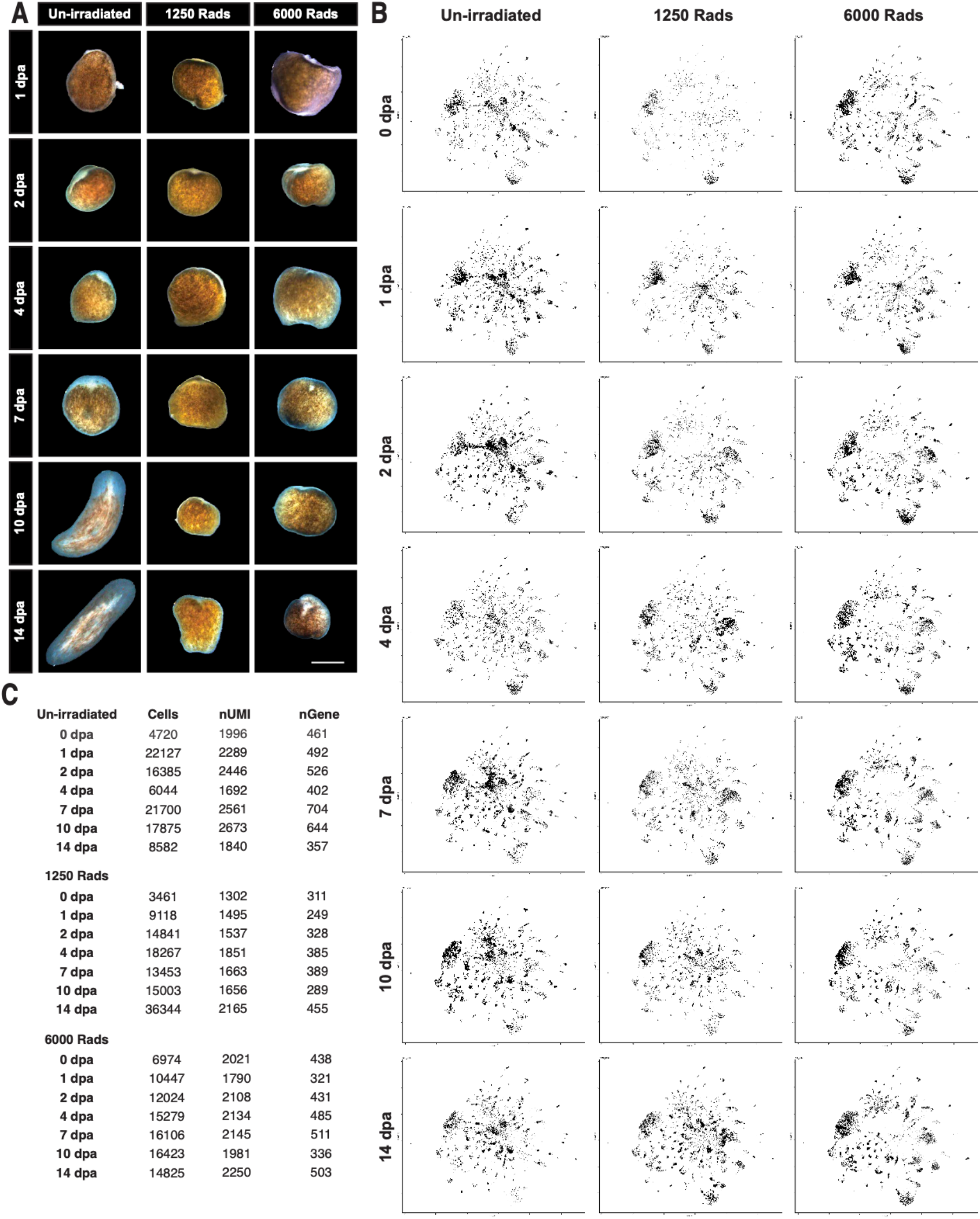
A single cell-reconstruction of planarian regeneration (A) Representative images of biopsies from un-irradiated, sub-lethally irradiated, and lethally irradiated animals imaged 1,2, 4, 7, 10, and 14 days post amputation. (B) UMAP embeddings cells sampled from each of the 21 conditions (see materials and methods for sub-sampling methodology) illustrating the change in tissue composition and captured transcriptional states across the dataset. (C) Number of cells captured, mean nUMI/cell, and mean nGene/cell for each of the 21 conditions. Scale = 500um.

**Extended Data Fig. 4.**
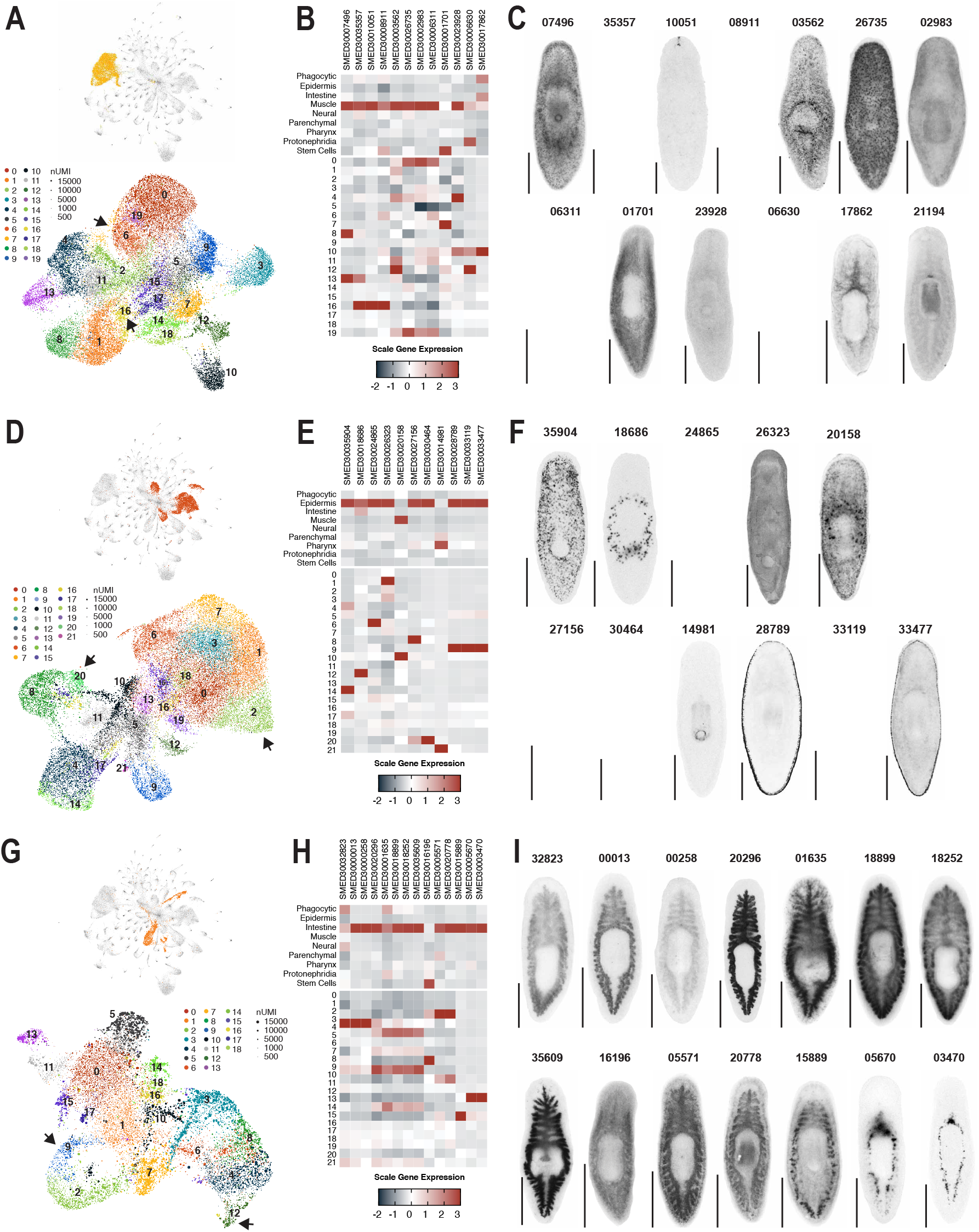
Selected marker genes used to confirm tissue annotation of muscle, epidermal, and intestinal clusters. UMAP embedding of global dataset with tissue highlighted and UMAP embedding of tissue cells colored by tissue sub-cluster ID for muscle (A), epidermis (B), or intestine (C). Scaled mean expression of cluster enriched genes by tissue and tissue sub-cluster for muscle (D), epidermis (E), and intestine (F) enriched genes. Whole mount in situ hybridization of tissue markers analyzed in D (G), E (H), and F (I).

**Extended Data Fig. 5.**
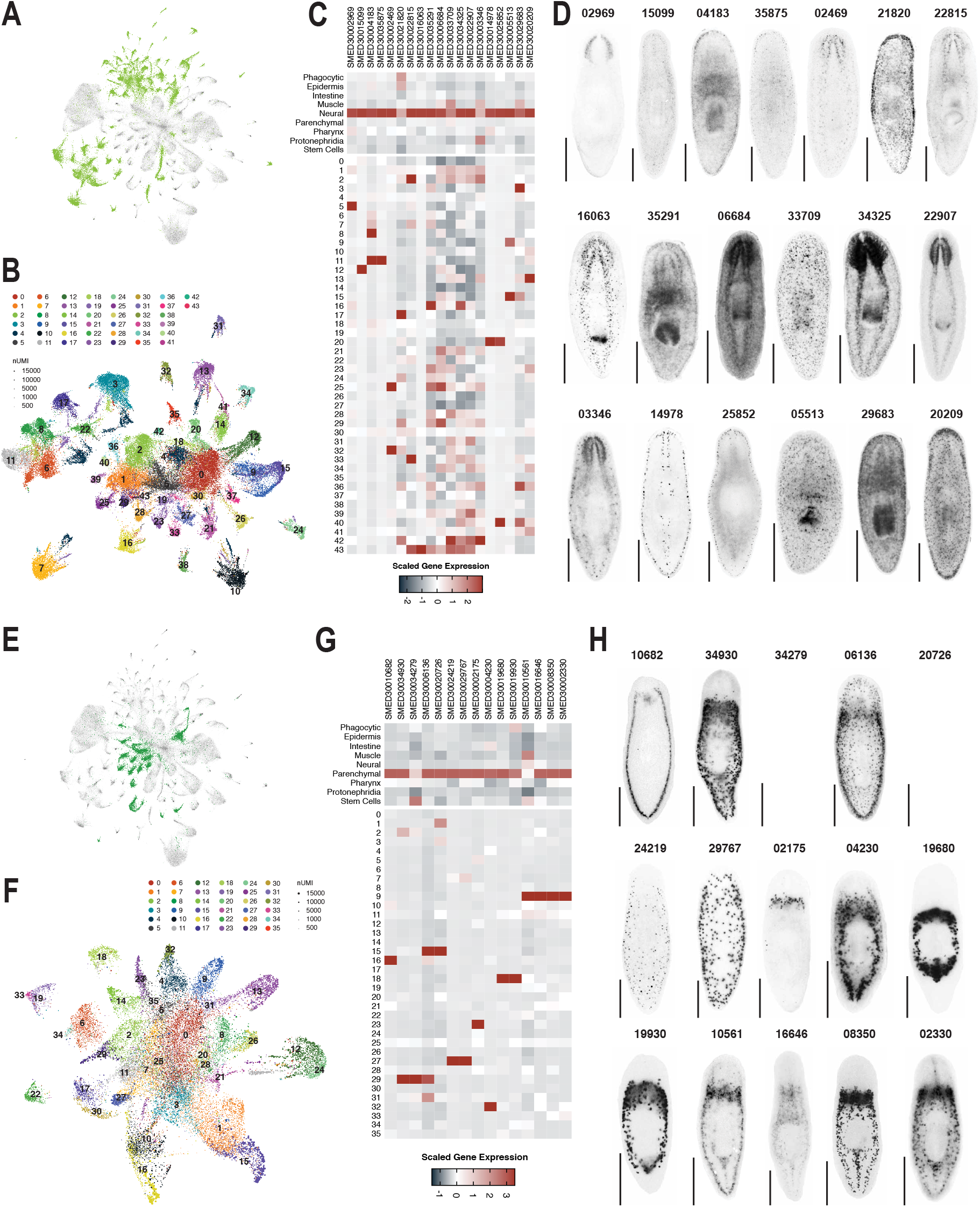
Selected marker genes used to confirm tissue annotation of neural and parenchymal clusters. UMAP embedding of global dataset with nervous system (A) or parenchyma (E) highlighted. UMAP embedding of neural (B) or parenchymal (F) cells colored by tissue subcluster ID. Scaled mean expression of cluster neural-enriched (C) or parenchymal-enriched (G) genes by tissue and tissue sub-cluster. Whole mount in situ hybridization of tissue markers analyzed in C (D) or G (H).

**Extended Data Fig. 6.**
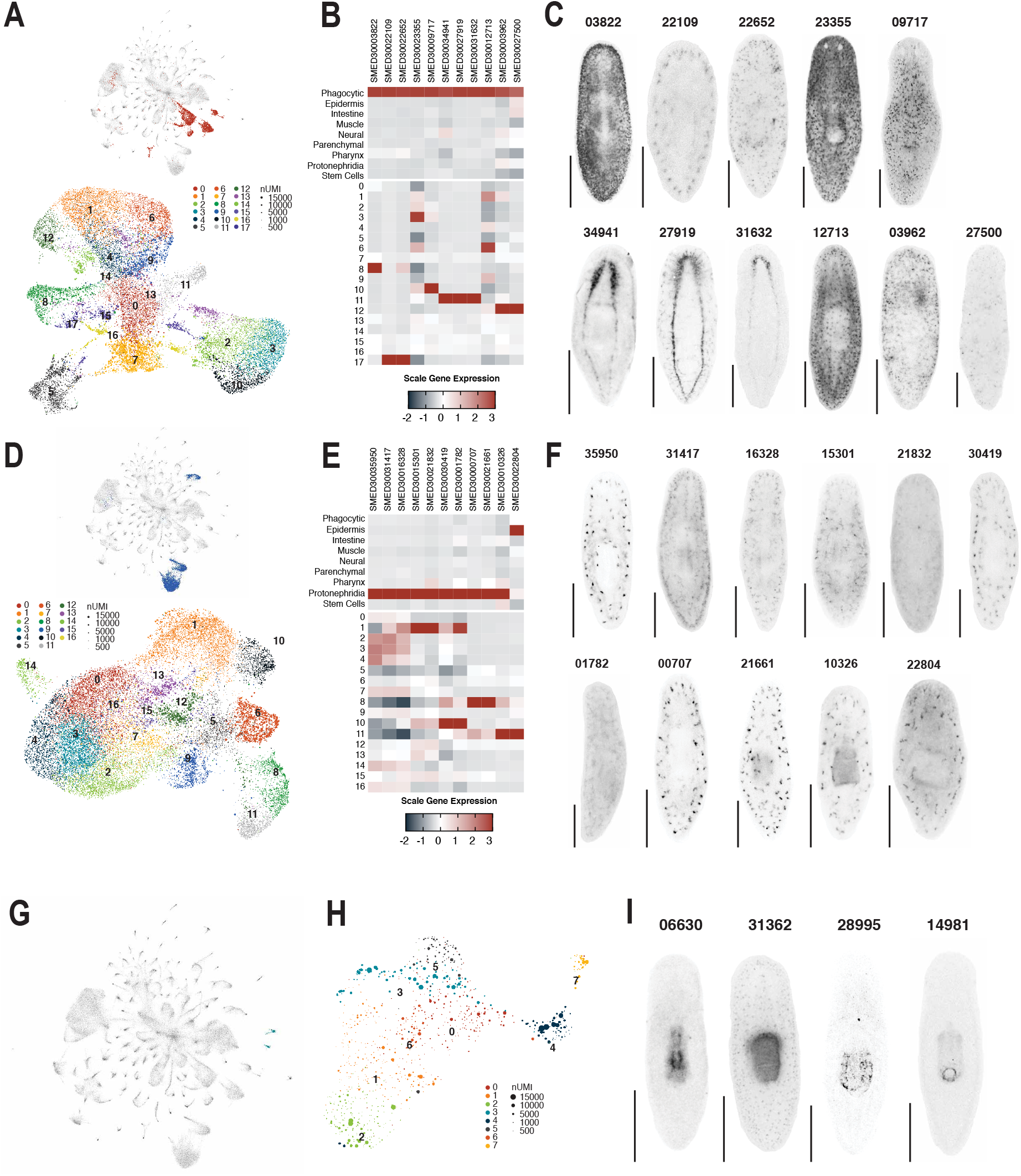
Selected marker genes used to confirm tissue annotation of phagocytic, protonephridial, and pharyngeal clusters. UMAP embedding of global dataset with tissue highlighted and UMAP embedding of tissue cells colored by tissue sub-cluster ID for phagocytic (A) and protonephridial (D) cells. Scaled mean expression of cluster enriched genes by tissue and tissue sub-cluster for phagocytic (D and protonephridia (F) enriched genes. Whole mount in situ hybridization of tissue markers analyzed in B (G) and E (F). (G) UMAP embedding of global dataset with pharyngeal clusters highlighted. (H) UMAP embedding of pharyngeal cells. (I) Whole mount in situ hybridization of pharynx-enriched genes.

**Extended Data Fig. 7.**
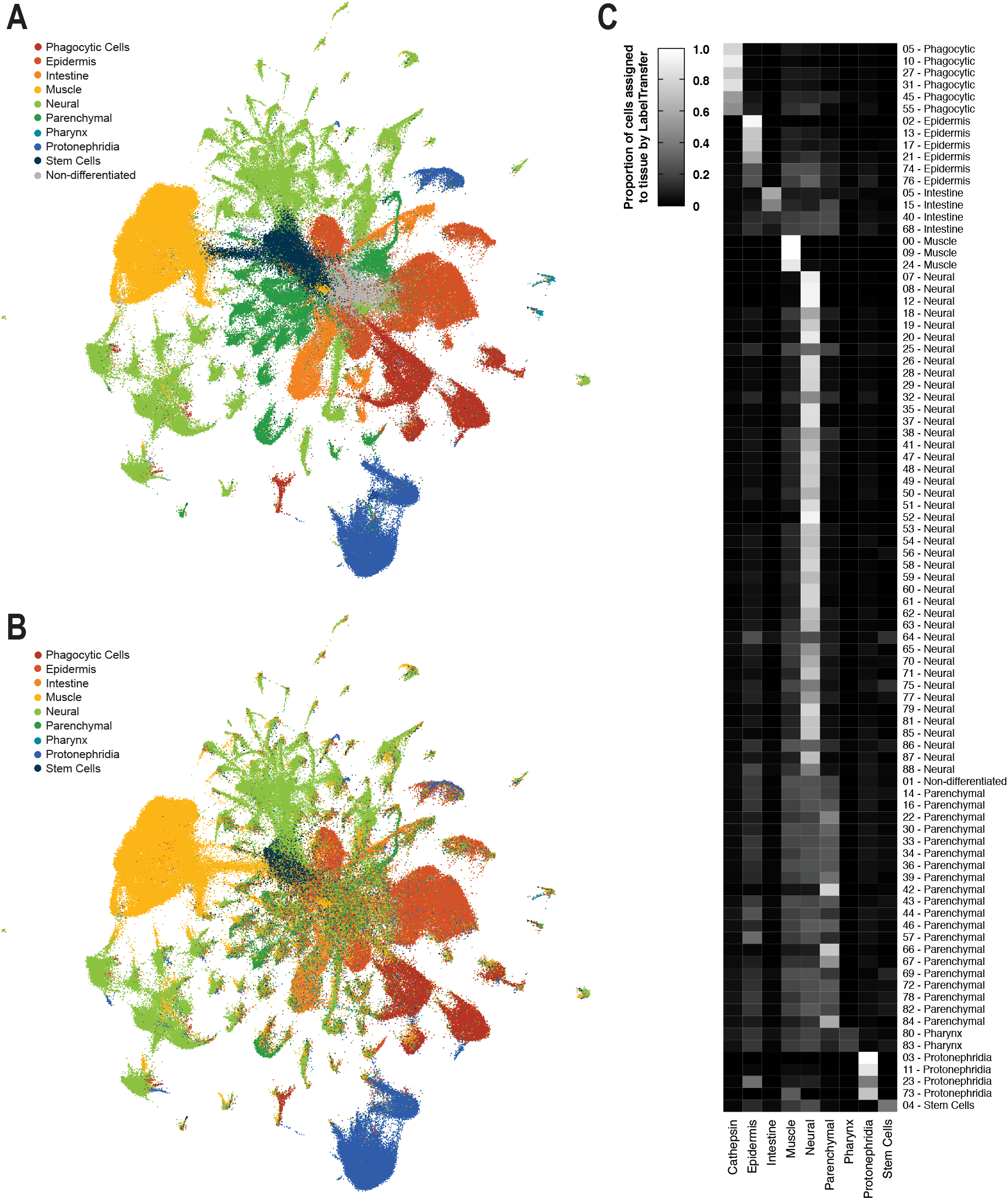
Confirmation of tissue annotation by TransferData. (A) Tissue annotation based on manual comparison of highly enriched genes to previously published single-cell planarian atlases (10,11). (B) Tissue annotation prediction made using Fincher et al. tissue annotations transferred to SPLiTseq dataset using Seurat’s TransferData function. (C) Proportion of cells from each global tissue cluster assigned to tissue lineages by TransferData.

**Extended Data Fig. 8.**
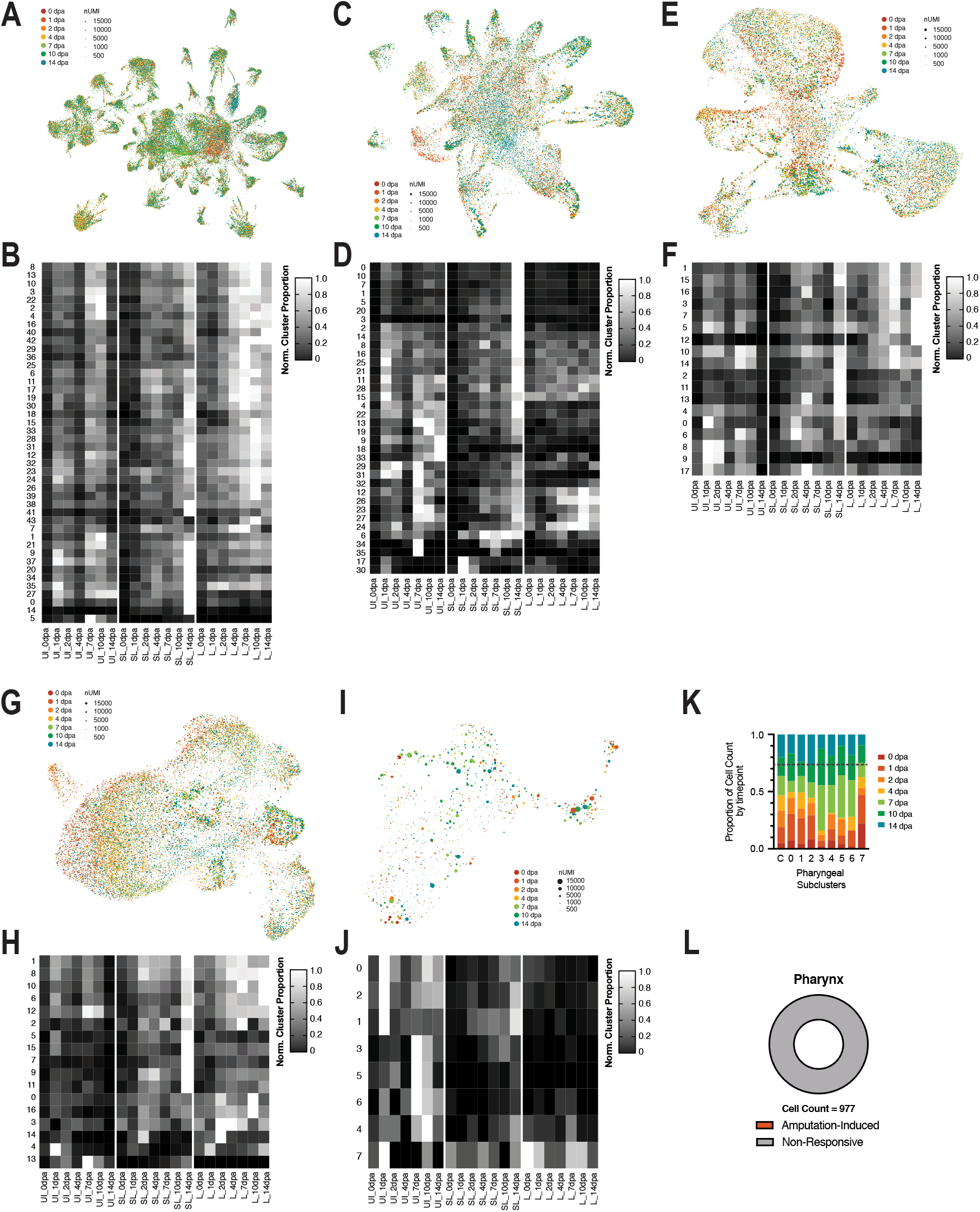
UMAP embedding of all neural (A), parenchymal (C), phagocytic (E), protonephridial (G), and pharyngeal (I) cells, colored by time. Scaled proportion of cells from each neural (B), parenchymal (D), phagocytic (F), protonephridial (H), or pharyngeal (J) sub cluster across sampled conditions, normalized to the sample in which the sub-cluster had maximum representation. (K) Proportion of cells from each pharyngeal sub-cluster that fall into each timepoint. (L) Proportion of pharyngeal cells that are in an amputation-specific sub-cluster.

**Extended Data Fig. 9.**
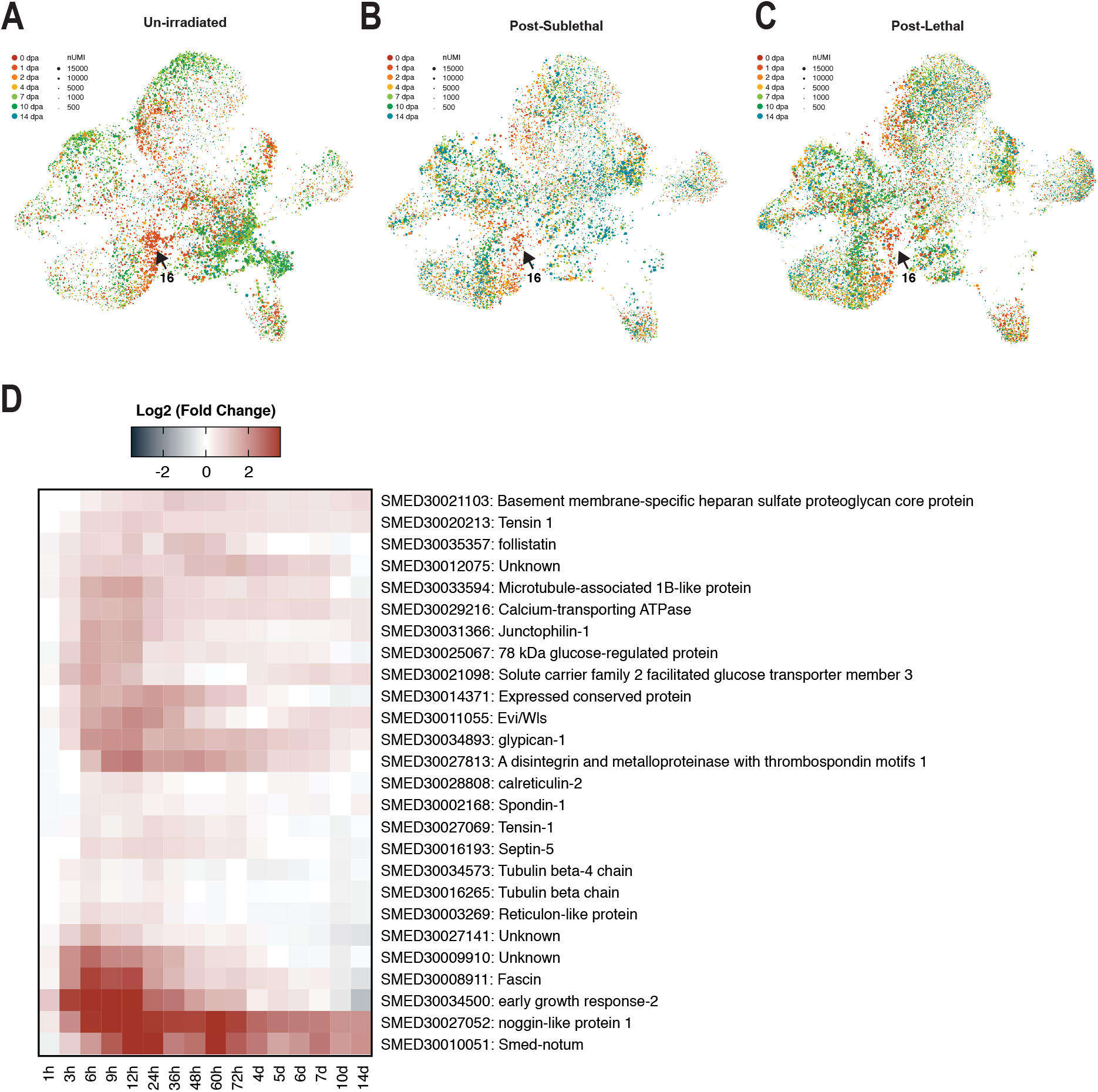
Muscle cluster tissue composition dynamics across treatments and wound-induced gene expression of screened genes. UMAP embedding of all muscle cells, colored by time after amputation and split by cells from biopsies taken from un-irradiated (A), sub-lethally irradiated (B), or lethally irradiated (C) animals. (D) Gene expression of screened muscle genes in bulk RNAseq dataset of planarian regeneration (*55*).

**Extended Data Fig. 10.**
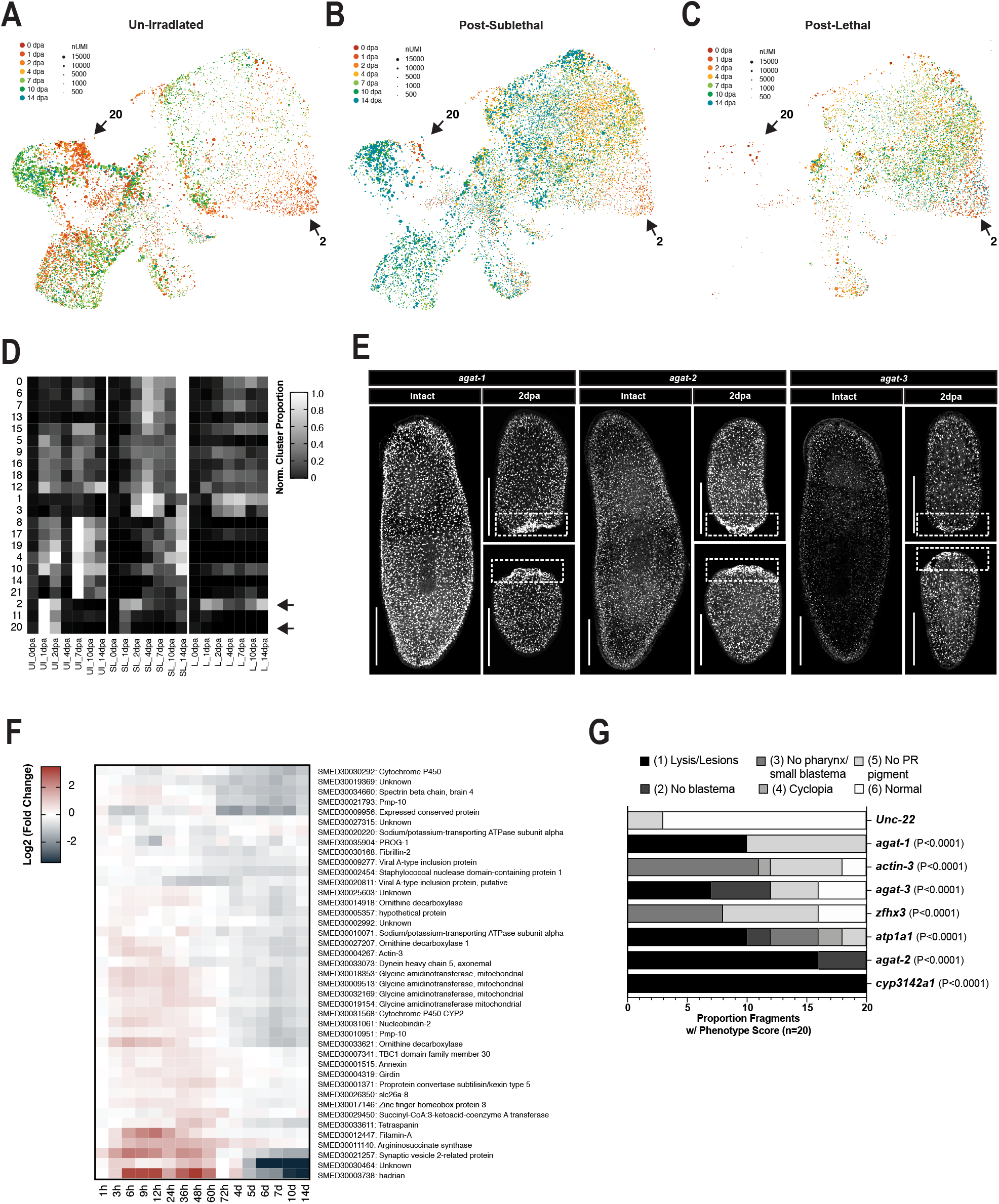
Additional data supporting wound-induced epidermal clusters and EC20-enriched genes requires for regeneration. UMAP embedding of all epidermal lineage cells, colored by time after amputation and split by cells from biopsies taken from un-irradiated (A), sub-lethally irradiated (B), or lethally irradiated (C) animals. (D) Scaled proportion of cells from each epithelial sub-cluster across sampled conditions, normalized to sample in which sub-cluster had maximum representation. (E) Maximum intensity projections from confocal stacks of whole mount fluorescent in situ hybridizations of EC20 enriched genes. (F) Gene expression of screened epidermal genes in bulk RNAseq dataset of planarian regeneration (*55*). (G) Scoring of phenotypes in RNAi treated animals in Figure 6G (n=20 posterior fragments). P values are Mann Whitney Test compared to Unc-22 Control. Scale = 500um.

**Extended Data Fig. 11.**
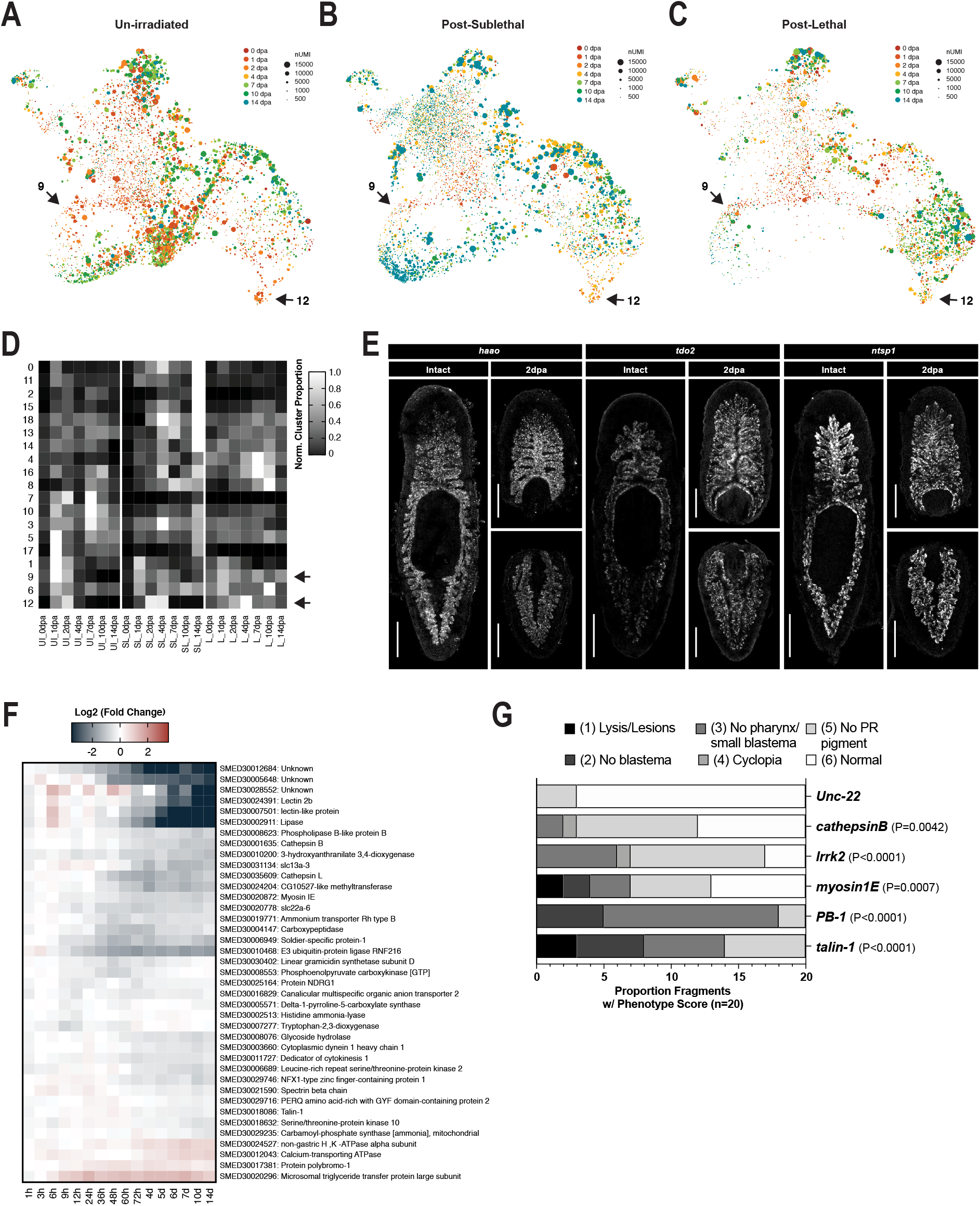
Additional data supporting wound-induced Intestinal clusters and intestinal genes requires for regeneration. UMAP embedding of all intestinal cells, colored by time after amputation and split by cells from biopsies taken from un-irradiated (A), sub-lethally irradiated (B), or lethally irradiated (C) animals. (D) Scaled proportion of cells from each epithelial sub-cluster across sampled conditions, normalized to sample in which sub-cluster had maximum representation. (E) Maximum intensity projections from confocal stacks of whole mount fluorescent in situ hybridizations of intestinal genes. (F) Gene expression of screened intestinal genes in bulk RNAseq dataset of planarian regeneration (*55*). (G) Scoring of phenotypes in RNAi treated animals in Figure 7I (n=20 posterior fragments). P values are Mann Whitney Test compared to Unc-22 Control. Scale = 500um.

## Supplementary Tables

**Supplementary Table 1. (excel file)**

Description of single cells and associated metadata

**Supplementary Table 2. (excel file)**

Cluster and sub-cluster enriched genes

**Supplementary Table 3. (excel file)**

Transcript IDs, annotation (if applicable), and RNAi phenotypes of all transcripts functionally tested in this study

